# A similarity matrix for preserving haplotype diversity among parents in genomic selection

**DOI:** 10.1101/2023.06.01.543227

**Authors:** Abdulraheem A. Musa, Norbert Reinsch

## Abstract

Mendelian sampling variability (MSV), determined by the heterozygosity and linkage phases of the parental haplotypes, quantifies the chance of producing offspring with high breeding values. Recent genomic selection criteria combine expected breeding values with MSV to maximize the chance of producing offspring with exceptional breeding values. These criteria, however, tend to select similar parents with high variability potential. Therefore, a measure of haplotype similarity is required to avoid this tendency and preserve diversity. Here, we derive this measure by pairing all potential gametes from two parents based on their segregating marker patterns. Subsequently, a similarity measure between two parents is defined as the absolute value of the covariance between the additive values of the paired gametes on a chromosome. A similarity matrix with absolute covariances as off-diagonal elements and MSVs as diagonal elements summarizes all pairwise similarities between parents. A parent’s similarity to itself equals its MSV. High similarity indicates that the parents share many heterozygous markers with large effects on a trait in the same linkage phase. The concept generalizes to multiple chromosomes, an aggregate genotype with multiple traits, and similarity between zygotes. We demonstrated the properties of the similarity matrix using empirical data. Through simulations, we showed that incorporating the matrix into genomic selection preserves up to 1630% more genetic variability and yields up to 7% more genetic gain relative to index selection in the long term. Although further research is needed, our results show that including similarity matrices preserves haplotype diversity and improves long-term genomic selection.

## Introduction

Owing to the prospect of accelerating genetic gain by shortening the generation interval and increasing the accuracy of estimated breeding values of young candidates, genomic selection (GS) has become a popular and essential tool for breeders (Meuwissen *et al*. 2001; Heffner *et al*. 2009; Hickey *et al*. 2017). In classical GS, the selection criterion is the genomic estimated breeding value (GEBV), computed as the sum of all the estimated additive effects of all alleles that make up an individual’s genotype (Meuwissen *et al*. 2001). The truncation selection (TS) of young candidates as parents based on their GEBV maximizes the population mean of the next generation. Its continued use, however, also promotes inbreeding as a side effect (Schaeffer 2006; Liu *et al*. 2015; Rutkoski *et al*. 2015; Lin *et al*. 2016). As a countermeasure, optimum contribution selection (OCS) (Meuwissen and Sonesson 1998; Sonesson *et al*. 2012; Woolliams *et al*. 2015) imposes constraints on the coancestry of selected parents, leading to more genetic progress in the long term.

Proposed selection criteria other than GEBVs promise more substantial genetic gain over a longer sequence of generations with recurrent selection. These criteria may also better preserve genetic variability in the breeding population. A group of these selection criteria focuses on the additive values (GEBVs) of predefined chromosomal sections rather than individuals to achieve highly favorable combinations of chromosomal segments following generations of repeated segregation and selection. As Kemper *et al*. (2012) presented, genotype building considers the best of two haplotype blocks in a group of diploid individuals. Individuals more likely to produce superior offspring when bred are selected in each generation. A related proposed method is the selection of parents based on optimal haploid values, which is the sum of maximal segmental GEBVs (Daetwyler *et al*. 2015). Similarly, when optimal population value selection (Goiffon *et al*. 2017) is applied, a set of parents is selected such that the best possible future offspring has the maximum additive value. Modification with a predefined fixed planning horizon is associated with look-ahead selection (Moeinizade *et al*. 2019), which maximizes the expected GEBV of the best progeny in the terminal generation.

The second group of proposed selection criteria also aims to provide an optimal trade-off between short- and long-term gains but does not use chromosome segmentation. Instead, they share the common principle of an index that is the sum of the expected GEBV of offspring plus a constant multiplied by the within-family standard deviation of the GEBVs of future offspring. As with expected GEBVs, these standard deviations are predicted from parental alleles and additive marker effects, either by simulating the segregation of parental haplotypes into gametes (Cole and VanRaden 2011; Bernardo 2014; Segelke *et al*. 2014; Mohammadi *et al*. 2015) or applying an analytical formula (Bonk *et al*. 2016; Santos *et al*. 2019). The usefulness criterion (Schnell and Utz 1976; Lehermeier *et al*. 2017) and expected maximum haploid breeding value (Müller *et al*. 2018) are examples of plant breeding with doubled haploid lines as offspring. Bijma *et al*. (2020) investigated the TS of parents based on various indices and found greater genetic gain in simulated diploid animals than for the conventional GS on GEBVs. In general, the constant in the standard deviation terms of these indices can be interpreted as selection intensities among future offspring, obtained either as an integral over the upper tail of the standard normal distribution or from order statistics (Müller *et al*. 2018) if only a limited number of offspring are considered.

As part of such indices, the within-family variation (Mendelian sampling variance, MSV) of additive genetic values quantifies the chance of obtaining offspring with high breeding values (Segelke *et al*. 2014; Bonk *et al*. 2016; Müller *et al*. 2018; Bijma *et al*. 2020). Therefore, parents with heterozygous genotypes at marker loci with large positive effects in coupling are more likely to be selected. However, breeders may be interested in keeping wide varieties of haplotypes in play over succeeding generations to avoid unintentional losses of genetic variability and to allow new favorable haplotypes to emerge through recombination in later generations. The latter is essentially the same aim ingrained in the first group of selection criteria mentioned above by design. Combinations of GEBV and Mendelian standard deviation, on the other hand, lack this inherent mechanism.

As a solution, we propose a similarity matrix among parents in which off-diagonal elements reflect pairwise haplotype similarities and diagonal elements reflect a parent’s similarity with itself, which equals the MSV of its gametes. Similar to how coancestry matrices are used to limit inbreeding in OCS, similarity matrices of this kind can be used to limit the selection of parents with similar haplotypes and preserve haplotype diversity.

In this paper, we draw upon previous results by Bonk *et al*. (2016) and introduce a computationally faster representation of their method for predicting the MSVs of gametes with the help of a covariance matrix of genotype indicators reflecting expected within-family linkage disequilibrium (LD). Second, we derive a measure of pairwise haplotype similarity for a single chromosome that relies on the same LD matrix. The concept is then extended to multiple chromosomes, multiple traits, and zygotes. Finally, we illustrate potential applications of the similarity measure for preserving haplotype diversity through simulation.

## Materials and Methods

In this section, we first review the method of Bonk *et al*. (2016) for calculating MSV and then present an equivalent but computationally faster representation. This representation will also be useful in defining our new similarity measure and its computation with large numbers of markers and as many individuals as needed for potential applications in an optimized selection or mating decisions.

### Mendelian sampling variance of gametes

An analytical approach for estimating the MSV of gametes was presented by Bonk *et al*. (2016). Previous attempts (e.g., Segelke *et al*. 2014; Mohammadi *et al*. 2015) estimated MSV through simulation. Both approaches use recombination rates, marker effects, and phased genotypes of parents as information and, thus, are equivalent, except for Monte Carlo errors. The exact formula for MSV is related to the breeding value *b*_*i*_ of individual *i* :

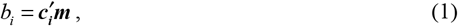

where 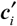 is a row vector of genotype indicators for biallelic markers (1, 0, and −1 for respective genotypes *AA, AB* or *BA*, and *BB* at each locus, where *A* is the reference allele whose effect is to be estimated) and ***m*** is a vector of additive marker effects of the population, most commonly allele substitution effects. Then, the MSV of the additive genetic values *g*_*i*_ of the gametes produced by a potential parent *i* can be computed, according to Bonk *et al*. (2016), as follows:

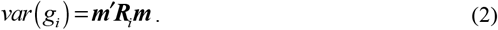

Here, ***R***_*i*_ is a parent-specific covariance matrix of additive genotype indicators in 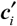 and is derived from the recombination rates and linkage phase of the parent *i*. Thus, the elements of ***R***_*i*_ reflect the within-family LD of markers. ***R***_*i*_ is a block-diagonal matrix with one block per chromosome. Using an appropriate mapping function, the elements *ρ*_*kl*_ of ***R***_*i*_ can be expressed in terms of genetic distances. Assuming Haldane’s mapping function (Haldane 1919), each element is calculated as:

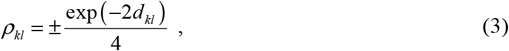

where the sign of off-diagonal elements depends on the linkage phase of markers and *d*_*kl*_ is the genetic distance between two markers in Morgan. The diagonal elements of ***R***_*i*_ (*d*_*kl*_ = 0) have a positive value of 0.25, while the off-diagonal elements corresponding to two markers on different chromosomes have a value of zero (*d*_*kl*_ =∞). All rows and columns of ***R***_*i*_ corresponding to homozygous markers are set to zero because they do not contribute to MSV.

The MSV of additive genetic values *g*_*ij*_ of zygotes from parents *i* and *j*, then, is the sum of the two independent contributions (Bonk *et al*. 2016) of the paternal and maternal gametes:

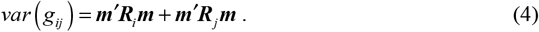

Note that in a scenario with only additive effects, the MSVs for additive genetic values remain unaffected if half the differences between homozygotes are used as additive marker effects instead of allele substitution effects (Bonk *et al*. 2016, page 10).

#### Equivalent representation

We propose computing the MSVs of gametes produced by parent *i* in a modified but equivalent manner as follows:

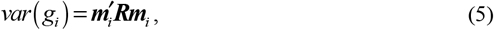

where ***m***_*i*_ is now a parent-specific vector of additive marker effects, incorporating the linkage phase of the markers via appropriately chosen marker effects signs, and matrix ***R*** is the expected within-family LD of markers, with elements as defined in ***R***_*i*_ but with all positive signs and without setting all rows and columns to zero for homozygous markers. For each marker *k*, a phase-indicator variable is set to *δ* _*ik*_ = 1 if the reference allele is observed on the first haplotype of individual *i* ; otherwise, it is set to *δ* _*ik*_ = −1. For any homozygous genotype, the indicator is set to zero *δ* _*ik*_ = 0. Then, each additive marker effect *m*_*k*_ is converted into an element *m*_*ik*_ by multiplying *m*_*k*_ by the respective indicator *δ* _*ik*_, *m*_*ik*_ = *δ* _*ik*_ *m*_*k*_. Thus, the sign of the respective additive marker effect is retained as in ***m*** in equation 2 if the reference allele is found on the first haplotype; otherwise, the sign is the opposite if the marker genotype is heterozygous. As homozygous loci do not contribute to MSV, they are assigned a zero value *m*_*ik*_ as an entry in ***m*** _*i*_. See Appendices 1 and 2 for an illustration and proof of the equivalence of equations (2) and (5).

Compared to adapting the signs of the off-diagonal elements of ***R***_*i*_ to the linkage phase of markers in a parent, which requires exploring all possible marker pairs, adapting vector ***m*** _*i*_ is easier because only single markers need to be explored. The expected within-family LD matrix ***R*** then has to be set up only once, making the computation of many MSVs faster than previously presented formulas (Bonk *et al*. 2016; Santos *et al*. 2019; De Abreu Santos *et al*. 2020). Further, in the following, the new representation will prove useful in deriving our similarity measure.

From the above definition, it is also immediately clear that a swap of haplotypes changes ***m***_*i*_ into −***m***_*i*_, i.e., the same vector with opposite signs. However, the resulting MSV remains unaffected, regardless of which one of two haplotypes is chosen as the first.

#### Multiple traits and the aggregate genotype

Breeders rarely evaluate and select parents based on a single trait but instead make decisions based on multiple traits—therefore, they are interested not only in the MSV for each trait but also their covariance. The MSV 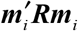 for a single trait of parent *i* extends to the vector ***g***_*i*_ of multiple traits,

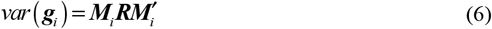

where each row of matrix ***M***_*i*_ is a vector 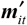 of additive marker effects for each trait *t* of the parent *i*. The diagonal elements of 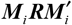 are the MSVs of all traits, whereas the off-diagonal elements are the Mendelian sampling covariances (MSCs) between traits (Bonk *et al*. 2016). Finally, by assuming a known vector of index weight ***a*** for all traits, the MSV for the aggregate genotype becomes

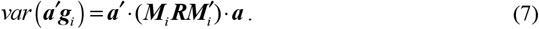

### Similarities between gametes from two parents

To derive our similarity measure, we consider two parents with a single chromosome and all the gametes they produce. First, we choose an arbitrary order of haplotypes from both parents and assign a segregation pattern to each possible gamete. Table 1 shows a demonstration using three biallelic markers and eight distinct segregation patterns. For example, pattern *A* − *B* − *A* implies that alleles from the first parental haplotype (*A*) are transmitted at the first and third loci while an allele from the second parental haplotype (*B*) is transmitted at the second locus. Recombination rates between adjacent markers (assumed to be *θ*_1,2_ = .1 and *θ*_2,3_ =.2) give the probabilities of gametes with a particular segregation pattern. For each of the first parent’s gametes, there is a matching gamete from the second parent with an identical segregation pattern and probability of occurrence. Additive values of matching gametes are, however, different and depend on which alleles (1 or 2) are present on the first (*A*) and second (*B*) haplotype of each parent.

**Table 1:** Segregation patterns (sequence of alleles from the first and second parental haplotypes), probabilities of occurrence, and genetic values of pairs of gametes produced by the first and second parents with matching recombination patterns.

| Segregation pattern/<br>Gametes produced | Probability | Genetic value |  |
| --- | --- | --- | --- |
|  |  | First parent | Second parent |
| $A - A - A$ | $\frac{1}{2} \cdot (1 - \theta_{1,2}) \cdot (1 - \theta_{2,3})$ | $\frac{1}{2} (m_1 + m_2 + m_3) = 0.61$ | $\frac{1}{2} (m_1 - m_2 + m_3) = 0.41$ |
| $A - A - B$ | $\frac{1}{2} \cdot (1 - \theta_{1,2}) \cdot \theta_{2,3}$ | $\frac{1}{2} (m_1 + m_2 - m_3) = 0.59$ | $\frac{1}{2} (m_1 - m_2 - m_3) = 0.39$ |
| $A - B - A$ | $\frac{1}{2} \cdot \theta_{1,2} \cdot \theta_{2,3}$ | $\frac{1}{2} (m_1 - m_2 + m_3) = 0.41$ | $\frac{1}{2} (m_1 + m_2 + m_3) = 0.61$ |
| $A - B - B$ | $\frac{1}{2} \cdot \theta_{1,2} \cdot (1 - \theta_{2,3})$ | $\frac{1}{2} (m_1 - m_2 - m_3) = 0.39$ | $\frac{1}{2} (m_1 + m_2 - m_3) = 0.59$ |
| $B - A - A$ | $\frac{1}{2} \cdot \theta_{1,2} \cdot (1 - \theta_{2,3})$ | $\frac{1}{2} (-m_1 + m_2 + m_3) = -0.39$ | $\frac{1}{2} (-m_1 - m_2 + m_3) = -0.59$ |
| $B - A - B$ | $\frac{1}{2} \cdot \theta_{1,2} \cdot \theta_{2,3}$ | $\frac{1}{2} (-m_1 + m_2 - m_3) = -0.41$ | $\frac{1}{2} (-m_1 - m_2 - m_3) = -0.61$ |
| $B - B - A$ | $\frac{1}{2} \cdot (1 - \theta_{1,2}) \cdot \theta_{2,3}$ | $\frac{1}{2} (-m_1 - m_2 + m_3) = -0.59$ | $\frac{1}{2} (-m_1 + m_2 + m_3) = -0.39$ |
| $B - B - B$ | $\frac{1}{2} \cdot (1 - \theta_{1,2}) \cdot (1 - \theta_{2,3})$ | $\frac{1}{2} (-m_1 - m_2 - m_3) = -0.61$ | $\frac{1}{2} (-m_1 + m_2 - m_3) = -0.41$ |

$A$ represents the alleles on the first haplotype, and $B$ represents the alleles on the second haplotype. The first parent has ordered haplotypes of $1-1-1/2-2-2$ , while the second parent has ordered haplotypes of $1-2-1/2-1-2$ . Recombination rates between adjacent markers are assumed to be $\theta_{1,2} = .1$ and $\theta_{2,3} = .2$ , and vector of population additive marker effects is assumed to be $\mathbf{m}' = [1 \quad .2 \quad .02]$ . Pattern $A - A - B$ , for example, has alleles $1-1-2$ in the first parent and $1-2-2$ in the second parent. Note that genetic value represents the value transmitted to the offspring.

In our example, the first parent has the ordered parental haplotype 1−1−1/ 2 − 2 − 2, while the second parent has the ordered parental haplotype 1− 2 −1/ 2 −1− 2. Consequently, the additive genetic value of the double recombinant *A* − *B* − *A* segregation pattern is either 0.41 or 0.61, depending on whether it descends from the first or second parent (Table 1). Obviously, the additive value assigned to a particular segregation pattern depends on the haplotype order. As mentioned above, the MSV for each parent does not change if haplotypes are swapped and, therefore, can unequivocally be derived from the probability distribution of segregation patterns.

The variance of the genetic values *g*_1_ from the gametes produced by the first parent in the example is calculated as follows:

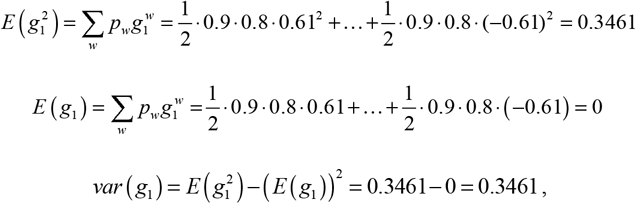

where *w* is the index of the segregation pattern, *p* is the probability of the segregation pattern *w*, and 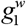 is the additive value of gamete transmitted from the first parent. Following the same procedure, the MSV for the second parent is calculated as 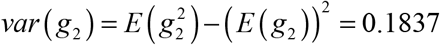.

Similarly, the probability distribution of segregation patterns allows us to compute the covariance between the additive genetic values of matching gametes from both parents. This covariance, however, is conditional on the chosen order of haplotypes. With *E* (*g*_1_) = *E* (*g*_2_) = 0, this covariance in our example is

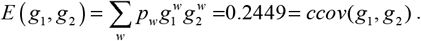

Enumerating all segregation patterns and probabilities of occurrence becomes difficult with hundreds to thousands of markers. However, using the previously mentioned equivalent matrix expressions, it becomes easy to compute the variances even when dealing with tens of thousands of markers (Bonk *et al*. 2016). Similarly, there is an equivalent matrix formula for the conditional covariance between the additive values of matching gametes. Relying on parent-specific vectors ***m***_*i*_ and ***m*** _*j*_, we define as the bilinear form

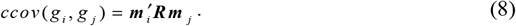

In our example (Table 1), the first haplotypes of both parents give rise to parent-specific vectors of marker effects 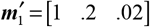 and 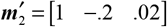 (see Additional File 1 (A.1-A.3) for numerical examples). Looking at the matrix expression, it becomes apparent that if both parents have markers with identical additive effects linked in the same manner on their first and second haplotypes, the MSVs in both families and the conditional covariance will be the same size. If turned into a correlation, the resulting value of one would then indicate the perfect equality of the ordered parental haplotypes and, hence, the equal distributions of their gametes’ additive values.

Considering a single chromosome, the sign of the conditional covariance changes if the haplotype order for one parent is swapped, turning either ***m***_*i*_ to −***m***_*i*_ or ***m*** _*j*_ to −***m*** _*j*_. Thus, the *ccov* (*g* _*i*_, *g* _*j*_) equals 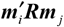 and 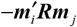 with equal probability if the order of haplotypes is randomly chosen. Consequently, the average conditional covariance over all possible haplotype orders is zero. The absolute value 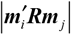 of our conditional covariance is independent of haplotype order. Therefore, a unique similarity measure valid for any possible order of parental haplotypes can be defined as

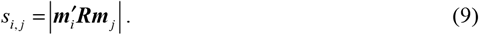

For a genome with several chromosomes, the similarity is calculated as the sum of absolute contributions from all chromosomes

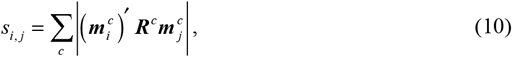

where ***R***^*c*^ is the diagonal block of ***R*** pertaining to chromosome *c*, and 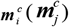 represents the parent-specific additive marker effects vector for parent *i* (*j*) on the same chromosome. An individual’s similarity to itself (i.e., *i* = *j*) equals that individual’s MSV. Therefore, it is possible to assemble a trait-specific similarity matrix ***S*** with MSVs for each parent on the diagonal and pairwise similarities *s*_*i,j*_ as off-diagonal elements for any set of individuals.

We emphasize that expectations are based on the univariate conditional distribution of segregation patterns rather than the bivariate joint distribution of gametes from two parents. The latter would result in a zero covariance because of the independence of the Mendelian sampling processes in different individuals.

#### Multiple traits and the aggregate genotype

When considering multiple traits, there is also a similarity between potential parents *i* and *j* with respect to the aggregate genotype. Each element *s*_*ij*_ of the respective similarity matrix is given by the following equation:

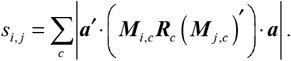

### Mendelian sampling variance of zygotes

#### An equivalent expression for MSV

We propose computing the MSV of progeny produced by parents *i* and *j* in the same way as described in Eq. (4):

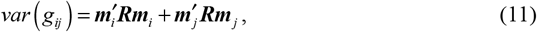

again, where ***R*** is the covariance matrix reflecting the expected within-family LD of markers, and ***m***_*i*_ (***m***_*j*_) represents the parent-specific additive marker effects vector for parent *i* (*j*).

#### MSV for multiple traits and the aggregate genotype

The extension to additive values for multiple traits (in vector ***g***_*ij*_) is presented by the Mendelian covariance matrix,

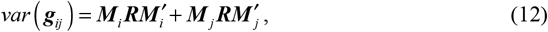

where each row ***m***_*it*_ and ***m***_*jt*_ of ***M***_*i*_ and ***M*** _*j*_, respectively, are vectors of the marker effects for trait *t* of parents *i* and *j*. Again, the result has zygotic MSVs for all traits on the diagonal and MSCs between traits as off-diagonals. Eq. (12) can be extended to multiple chromosomes as follows:

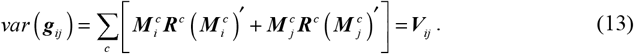

Since ***V***_*ij*_ is independent of the chosen order of haplotypes, the MSV for the aggregate genotype is unequivocally given by the following quadratic form:

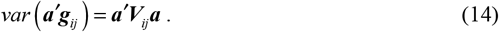

### Similarities between zygotes of two pairs of parents

A common breeding interest is determining the optimal parents for mating and the number of mates to assign to each parent, which can be determined using a gametic similarity matrix. However, in some cases, such as multiple ovulation and embryo transfer, determining the optimal parent pairs or the optimal number of offspring that particular parent pairs should produce is of interest. Consequently, a measure of the similarity between the genetic values of zygotes produced by two parent pairs can be useful. Similar to gametes, this measure can also be calculated from the probability distribution (see B.1 in File S1 for the derivation). This similarity can also be expressed in matrix notation, which is much more convenient for many markers. If ***m***_*i*_, ***m*** _*j*_, ***m***_*u*_ and ***m***_*v*_ are the marker effect vectors of parents *i, j, u*, and *v*, then the similarity between the additive genetic values (Mendelian sampling values) of zygotes produced by parent pairs *ij* and *uv* in the case of a single chromosome is:

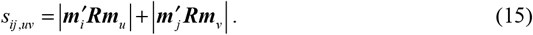

Eq. (15) operates independently of the order of haplotypes for all four parents. For multiple chromosomes, we maintain this independence by taking the sum of absolute values chromosome by chromosome and then summing them:

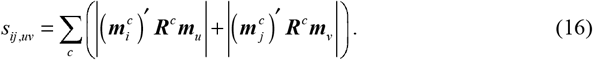

Finally, these similarities *s*_*ij,uv*_ define the off-diagonal elements of a similarity matrix ***S*** between pairs of parents (families) with respect to the makeup of the haplotypes in the zygotes of these families. The MSV of the respective family (which is equal to the similarity of a family with itself) represents the diagonal elements *s*_*ij,ij*_. Numerical examples of the derivation of MSV and the similarity between the zygotes of parent pairs *ij* and *uv* are also provided in File S1 (B.1).

#### Similarities for multiple traits and the aggregate genotype

For two pairs of parents *ij* and *uv*, the similarity measure *s*_*ij,uv*_ for the aggregate genotype, which does not depend on the haplotype order of any of the four involved parents _*i*_, *j, u*, and *v*, becomes

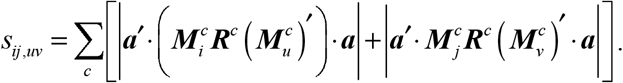

Then, the MSVs of the aggregate genotype define the diagonal elements, and the similarities *s*_*ij,uv*_ define the off-diagonal elements of the similarity matrix ***S*** between pairs of parents. See Appendix 3 for the extension of the similarity measure to monoecious species and the order of parents in dioecious organisms.

### Standardized similarity

The similarity matrix ***S*** of either gametes or zygotes can be standardized by pre- and post-multiplying with a diagonal matrix ***D***^−1^ with inverse Mendelian standard deviations as diagonal elements: (17)

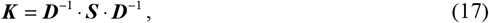

where 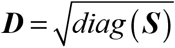. Thus, all diagonal elements of ***K*** are equal to 1, and all off-diagonal elements are strictly non-negative and lie within the range of 0≤ *k*_*ij*_ ≤1.

### Potential application of the similarity matrices for optimum mate allocation

The similarity matrices may be used to optimize mate allocation (OMA) such that optimal allocations of parents ***n***_*t*_ in generation *t* are obtained by maximizing the average expected genetic return 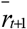 in the next generation under a given constraint on the average haplotype similarity 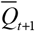. The model can be written as follows:

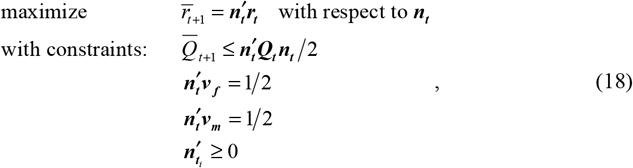

where vector ***r***_***t***_ denotes the selection criterion, such as breeding values or indices that combine breeding values and MSVs for parents in generation *t*. ***Q***_***t***_ is a similarity matrix that denotes ***S***_***t***_ or ***K***_***t***_ for gametes, ***v*** _***f***_ and ***v***_***m***_ are vectors of indicators (0/1) for females and males. This optimization strategy can be extended to parent pairs by deriving ***r***_***t***_ and ***Q***_***t***_ for parent pairs and by defining the genetic contributions of the parent pairs to account for user-specific constraints, such as the maximum/minimum allocations of parent pairs.

Although the similarity matrix ***S*** is positive semidefinite in theory, it may be indefinite in practice, for instance, due to rank deficiency or numerical inaccuracy. Then the similarity matrix can be approximated to the nearest positive definite matrix. The linear optimization problem with quadratic constraints can be solved using some efficient algorithm to obtain the optimal allocations of parents ***n***_*t*_ for any given data.

### Empirical Data

We analyzed an empirical dataset of five paternal half-sib families of Holstein-Friesian cows from Hampel *et al*. (2018) to demonstrate our approach. The publicly available dataset contains 265 individuals (32–106 candidates per paternal half-sib family), pedigree information, genotypic data, and a physical map. We approximated genetic map positions linearly by converting the physical map positions in Mbp into cM. The genotypic data contains 39,780 markers on the autosomes. The marker genotypes were phased with *hsphase* version 2.0.2 (Ferdosi *et al*. 2014), and unknown phases were removed, leaving 10,304 markers. The marker effects of this population were estimated in a previous study (Melzer *et al*. 2013).

We used all 265 individuals to derive similarity matrices using the gametic approach. To demonstrate their use for visualizing the haplotype diversity of a population, we graphically displayed these matrices using *corrplot* version 0.92 (Wei and Viliam 2021) and performed family-wise clustering using *pheatmap* version 1.0.12 (Kolde 2019).

### Simulated Data

To demonstrate the use of the similarity matrix for hedging haplotype diversity, we simulated a base population of 1,000 cattle (500 males and 500 females) with 10 chromosomes using *AlphaSimR* version 1.3.4 (Gaynor *et al*. 2021) and the default genomic and demographic parameters for cattle (MacLeod *et al*. 2013). We assumed a sex-averaged genetic map with an average length of 1 M per chromosome. The genome contains 10,000 single nucleotide polymorphisms (SNPs) and 2,000 QTLs, with 1,000 SNPs and 200 QTLs on each chromosome. We simulated additive QTL effects for a complex trait using a normal distribution with a mean of 0 and a variance of 100. The QTL effects were summed to obtain true breeding values (BVs) for all individuals, and trait heritability of 0.25 was simulated by adding residual effects to the BVs.

Then, we selected and mated parents based on various selection schemes (Table 2) for 50 generations, with each simulation run starting from the same base population. The selection schemes were created by combining selection criteria (BV and index) and rules (TS and OMA).

**Table 2:**
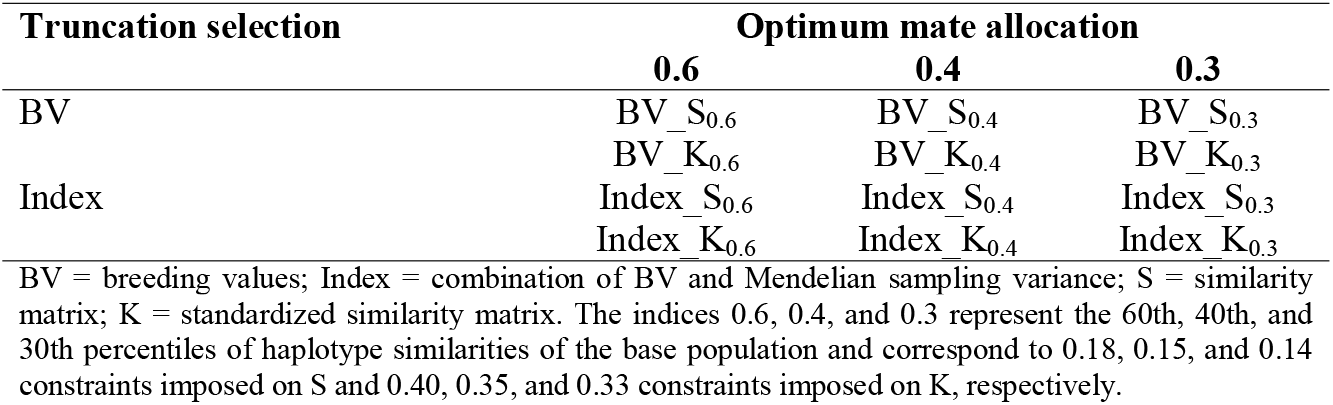
Overview of all selection schemes.

| Truncation selection | Optimum mate allocation |  |  |
| --- | --- | --- | --- |
|  | 0.6 | 0.4 | 0.3 |
| BV | BV_S<sub>0.6</sub> | BV_S<sub>0.4</sub> | BV_S<sub>0.3</sub> |
|  | BV_K<sub>0.6</sub> | BV_K<sub>0.4</sub> | BV_K<sub>0.3</sub> |
| Index | Index_S<sub>0.6</sub> | Index_S<sub>0.4</sub> | Index_S<sub>0.3</sub> |
|  | Index_K<sub>0.6</sub> | Index_K<sub>0.4</sub> | Index_K<sub>0.3</sub> |

BV = breeding values; Index = combination of BV and Mendelian sampling variance; S = similarity matrix; K = standardized similarity matrix. The indices 0.6, 0.4, and 0.3 represent the 60th, 40th, and 30th percentiles of haplotype similarities of the base population and correspond to 0.18, 0.15, and 0.14 constraints imposed on S and 0.40, 0.35, and 0.33 constraints imposed on K, respectively.

### Selection criteria

Based on the assumption of known additive marker effects, an average genetic map, and phased genotypes, we calculated BV *b*_i_ for parent *i* using Eq. (1), and calculated the index according to Bijma *et al*. (2020) as follows:

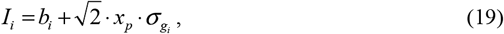

where *x*_*p*_ is the standardized truncation point belonging to the selected proportion *p* and 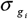 is the Mendelian standard deviation (square root of MSV) of the additive genetic values of gametes produced by parent *i*.

### Selection rules

For TS, the top 1% (5) of males and 50% (250) of females were selected according to selection criteria and mated randomly. The females were allowed one mating, while the males were assigned 50 females. Each mating produced four offspring, keeping the population size constant at 1,000 individuals. To ensure an equal number of male and female offspring, sexes were systematically assigned (i.e., male, female, male, etc.).

For OMA, we derived ***S***_*t*_ for all potential parents in each generation *t* using the gametic approach and scaled it using its maximum element (or MSV) to have a consistent constraint across generations. Additionally, the matrix was standardized to obtain ***K***_*t*_. When ***S***_*t*_ was not positive definite, we used the *nearPD* function in *Matrix* version 1.5-4 (Douglas and Maechler 2023) to approximate positive definiteness. The *nearPD* algorithm adapts the modified alternating projections method of Higham (2002) and adds procedures to guarantee positive definiteness (Maechler 2022).

Then the optimum allocations of parents ***n***_*t*_ to the next generation were derived at varying constraints on average haplotype similarity 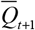 (see Table 2) by solving the optimization problem in Eq. (18) using *gurobi* package version 10.0-0 (Gurobi Optimization and LCC 2022). The constraints were selected based on the 60%, 40%, and 30% quantiles of the haplotype similarities of the base population.

Since females have limited offspring, the top 50% (250) of females were also selected based on either BVs or indices, and their allocations were forced to be equal, while unselected females received no allocations. Therefore, only the males were optimized, with their allocation limited to a maximum ratio of one male to 50 females to match scenarios with TS on BVs and indices. Thus, the minimum number of eligible males for mating was 5. The remaining conditions were identical to those in the selection on BVs or indices scenarios, except that males were randomly assigned to females based on the probabilities of their allocations.

### Parameters monitored

Several variables were monitored across the selection schemes and averaged over all 100 repetitions, which served as a basis for comparing the performance of the selection schemes. The first monitored variable is genetic gain (genetic standard deviation units), calculated as the mean increase in breeding value relative to the base population divided by the genetic standard deviation of the base population. The second monitored variable is the genetic variance expressed as the standard deviation of BVs for all 1,000 offspring. Pedigree-based inbreeding in each generation was also monitored and obtained using the *PedInbreeding* function of *optiSel* package version 2.0.6 (Wellmann 2023). Lastly, we monitored the number of favorable QTL alleles lost, the mean favorable QTL allele frequency, and the number of SNPs alleles lost.

All statistical analyses were performed using programs written by the authors and packages in R version 4.2.1 (R Core Team 2022).

## Results

We implemented a computationally faster approach for estimating MSV, building on the method described by Bonk *et al*. (2016). We benchmarked the time required to compute MSVs for 265 parents with 10,304 markers for a single trait (File S2) and found our approach 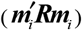 was 73 times faster than the previous approach (***m***′***R***_*i*_***m***, Bonk *et al*. 2016). Furthermore, the use of parent-specific ***m***_*i*_ vectors was essential for measuring the similarity among the Mendelian sampling values of individuals.

Considering the example data from Hampel *et al*. (2018), the derived genomic relationship matrix (GRM) (VanRaden 2008) using marker genotypes clearly showed the family structure in the population, with the progeny within a family being more similar to each other than to that of other families (Figure 1). However, this observed structure/pattern was not pronounced in the gametic similarity matrices because they mirror the Mendelian segregation patterns of the individuals (Figure 1A: milk fat, protein, and pH) rather than marker genotypes as in the GRM.

**Figure 1.**
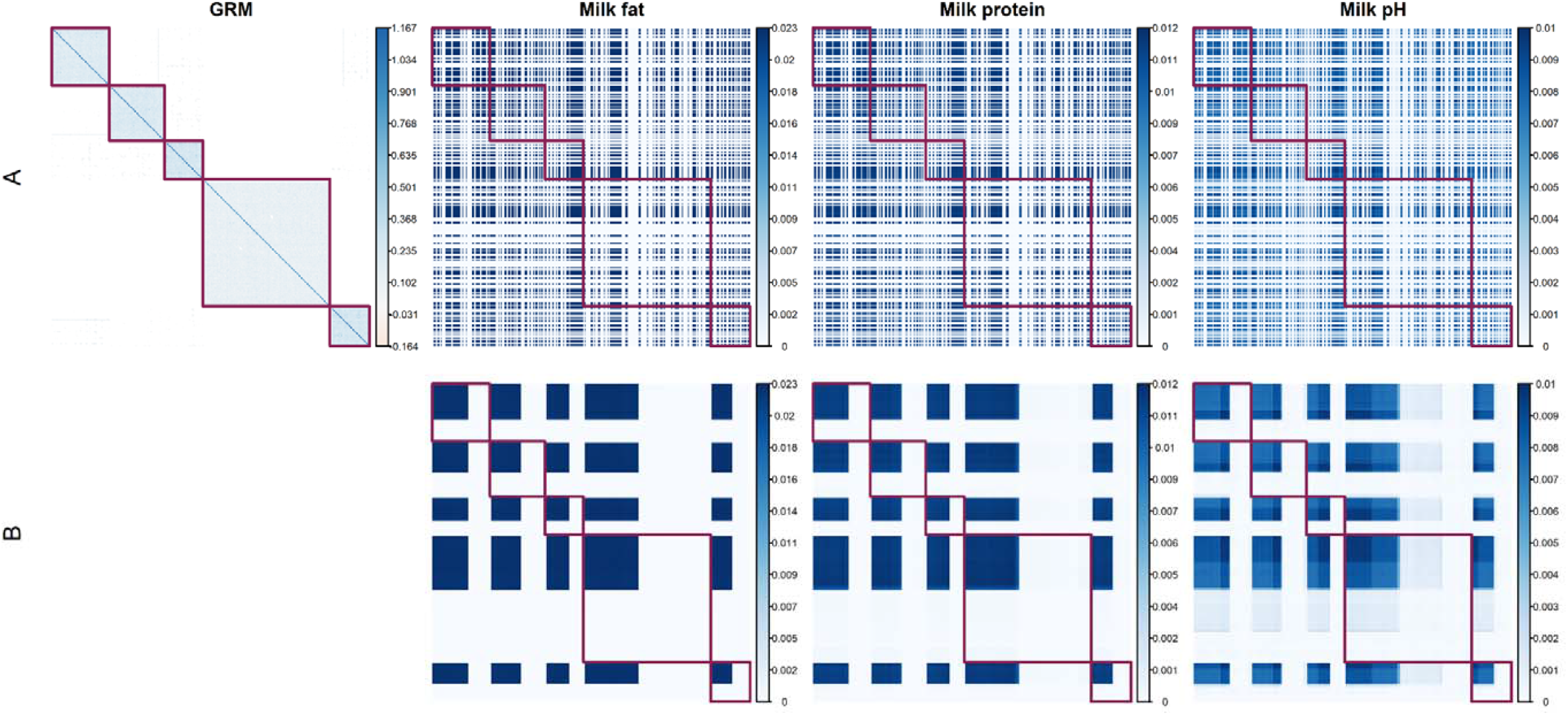
Genomic relationship matrix (GRM) and similarity matrices for all chromosomes (A) and Euclidean clustering of similarity matrices for all chromosomes (B) for milk fat, protein, and pH. The red blocks demarcate each paternal half-sib family. Parents are arranged according to their pedigree.

The diagonal elements of each matrix were the MSV of each parent, and the off-diagonal elements were the similarities among the Mendelian sampling values of potential parents. Thus, the values of these matrices depended on the linkage phase, heterozygosity, genetic architecture of traits (marker effect size), and distance between markers on a chromosome. Parents with common heterozygosity of markers with large effects for a trait in the same linkage phase appeared to be more similar than those with small effects. The latter appeared to be more similar than those that were homozygous at those loci (see numerical examples in A.3 File S1). Consequently, individuals with comparable MSV values due to common heterozygous loci appeared similar regardless of their family.

Clustering the individuals within a family revealed pronounced similarity structures across families (Figure 1B). The similarity matrices differed across traits, with pairwise similarities ranging from 0 to 0.023 in milk fat, 0 to 0.012 in milk protein, and 0 to 0.01 in milk pH. However, patterns of similarity encompassing all families were consistently observed for all traits. Standardized similarity matrices (Figure 2) revealed a similar trend to that depicted in Figure 1 but highlighted small similarities better, with pairwise similarities ranging from 0.007 to 0.9997 in milk fat, 0.017 to 0.998 in protein, and 0.020 to 0.997 in pH.

**Figure 2.**
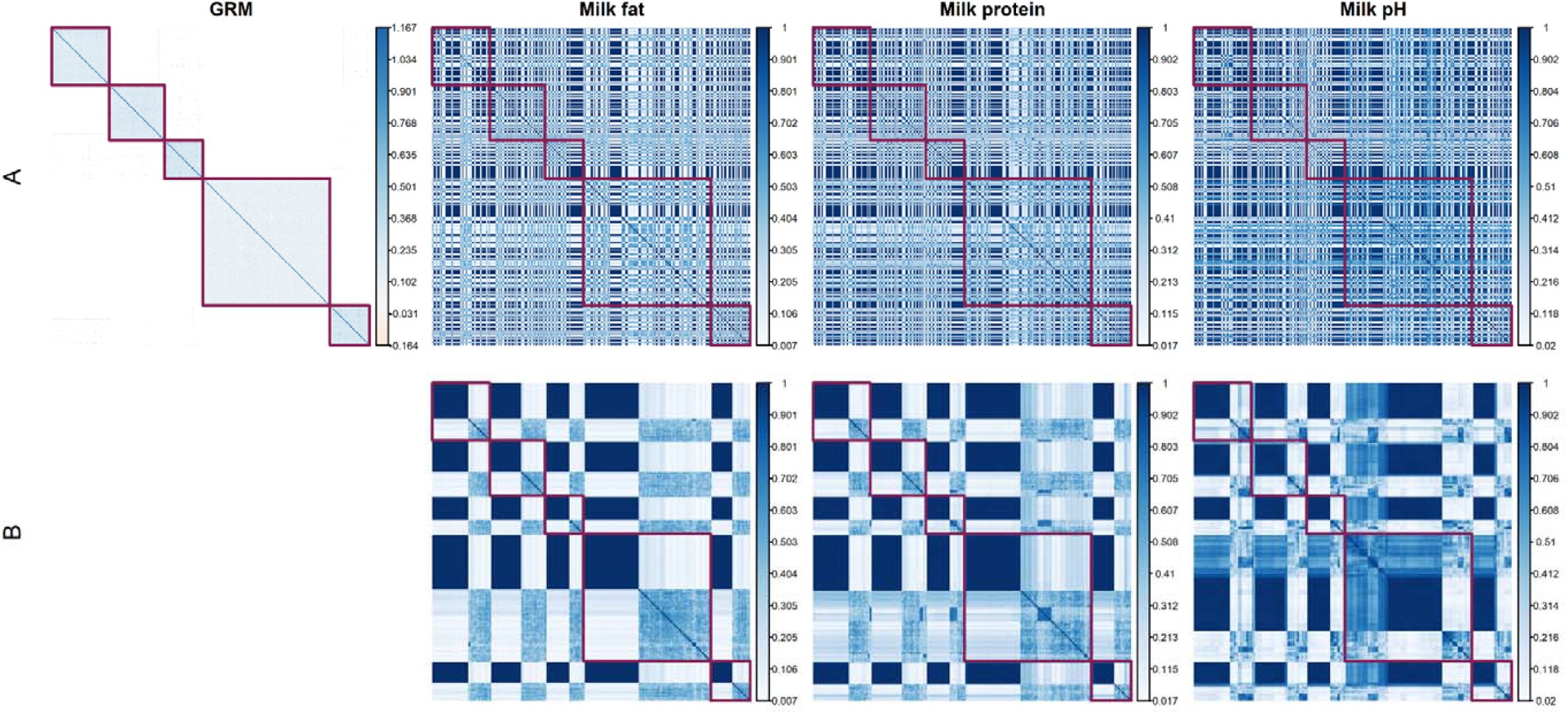
Genomic relationship matrix (GRM) and standardized similarity matrices (A) and Euclidean clustering of standardized similarity matrices (B) for milk fat, protein, and pH. The red blocks demarcate each paternal half-sib family. Parents are arranged according to their pedigree.

The effects of heterozygosity and trait genetic architecture were observed in chromosome 4, where individuals in family 4 appeared to be more variable than those in family 3 regarding milk fat, whereas the opposite was observed for milk protein (Figure S4 in File S3). Besides the apparent effect of various traits on the structure of the matrices, comparing the matrices from different chromosomes further showed the effect of genetic architecture (File S3; Figures S4 and S5). In milk fat, for example, the pairwise similarities ranged from 0 to 0.023 on chromosome 14, whereas the values on chromosome 4 were approximately zero (Figure S4 in File S3). Ignoring the effect of heterozygosity, such a result is expected if a chromosome (here: 14) has either many markers with small effects or few markers with large effects on milk fat than the other chromosome (here: 4).

The effects of optimizing mate allocation using the similarity matrix on genetic gain and variance over the long term are depicted in Figure 3. For brevity, the results presented pertain only to OMA schemes involving the index since those involving BV exhibit the same trend (Figure S6 in File S3). Imposing a small constraint on haplotype similarity (Index_S_0.6_ scenario) resulted in a greater long-term genetic gain at the expense of a slightly lower short-term genetic gain than their TS counterparts, whereas imposing large constraints (Index_S_0.4_ and Index_S_0.3_) resulted in less genetic gain across generations. The larger the constraint imposed on haplotype similarity, the smaller the short-term genetic gain and the greater the potential for long-term genetic gain. In this simulation, the TS of parents on the index produced marginally lower short-term genetic gain but 3% greater long-term genetic gain than the TS on BV (Figure 3A). Index_S_0.3_ produced 11% less genetic gain than TS on the index, while Index_S_0.6_ produced 2% more in the terminal generation. However, the long-term genetic variability preserved by the OMA schemes was consistently higher than their TS counterparts (Figure 3B). While the TS on the index preserved 41% more genetic variability than BV, the OMA schemes further amplified this effect, with Index_S_0.6_ preserving 74% and Index_S_0.3_ preserving 312% more genetic variability than the index, though at the expense of lower short-term genetic variance.

**Figure 3.**
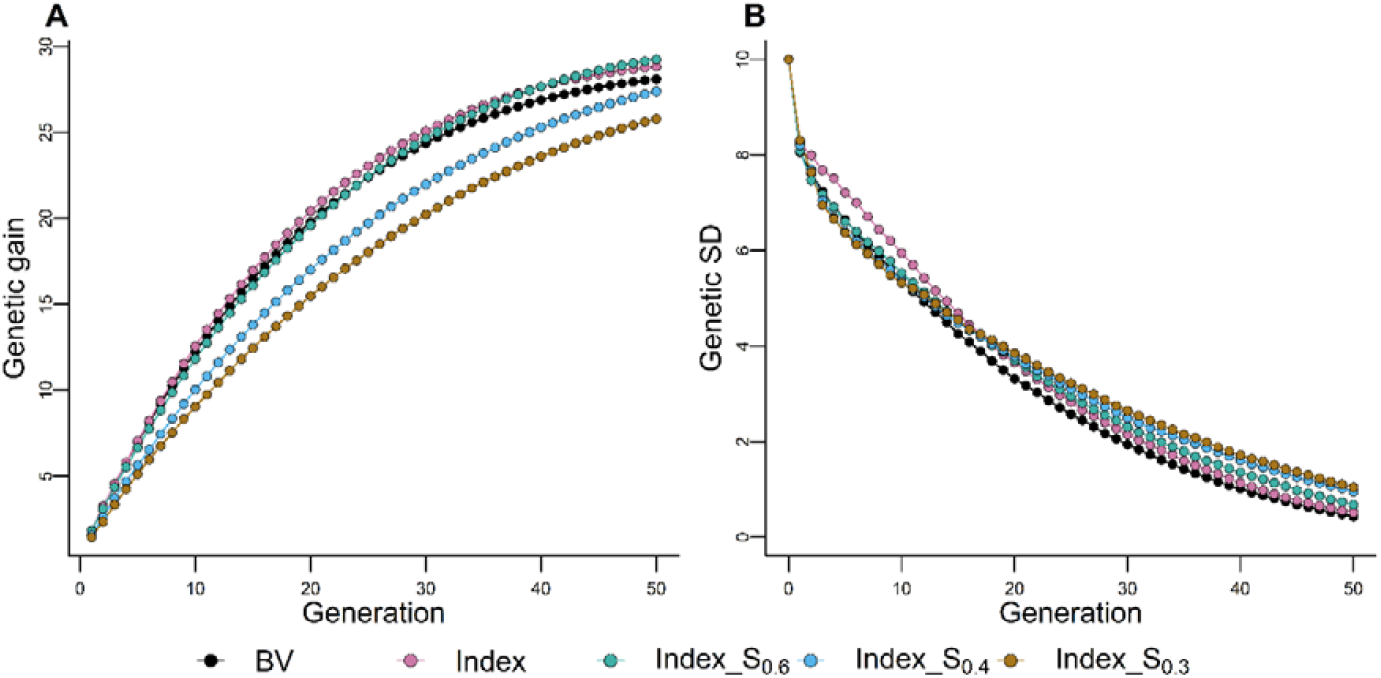
Effect of similarity matrix (S) on the cumulative genetic gain in genetic standard deviation (A) and genetic standard deviation (B). The selection schemes Index_S_0.6_(_0.4_,_0.3_) optimize mate allocation by maximizing the index combining breeding value (BV) and Mendelian sampling variance under various constraints (0.6, 0.4, and 0.3) on the haplotype similarity of parents. The results maximizing the BV are presented in Figure S6 in File S3. Results are reported for 100 simulation runs.

On the other hand, the performance of OMA schemes involving the standardized similarity matrix had a much greater impact on long-term genetic gain than those involving the unstandardized matrix (Figures 4 and S7 in File S3), yielding up to 5 – 7% more genetic gain than the index in the terminal generation (Figure 4A). OMA schemes using the standardized similarity matrix preserved between 179 – 1630% more genetic variability than those utilizing the TS on the index (Figure 4B) but sometimes produced lower short-term genetic variances, though not as severe as those involving unstandardized similarity matrices (see Figure 3B). These results show that OMA schemes using the standardized similarity matrix significantly preserves genetic variability without negatively affecting genetic gain in the long term.

**Figure 4.**
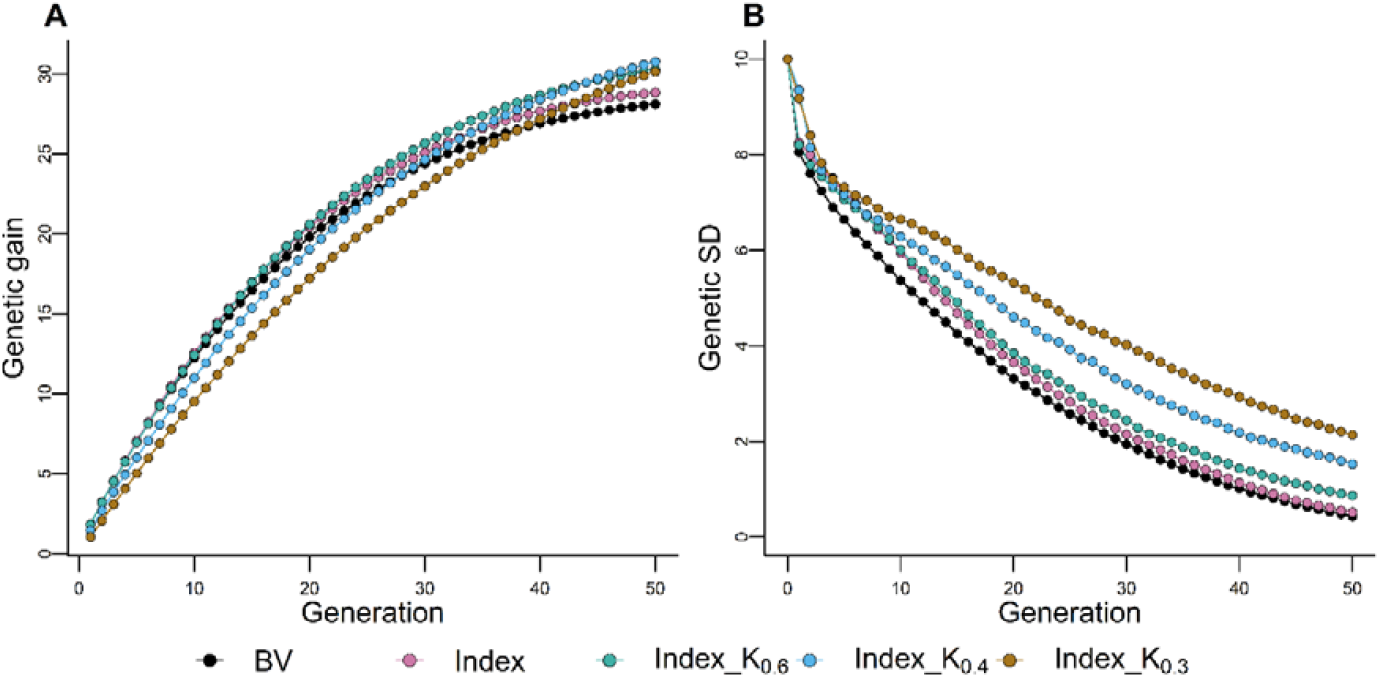
Effect of standardized similarity matrix (K) on the cumulative genetic gain in genetic standard deviation (A) and genetic standard deviation (B). The selection schemes Index_S_0.6_(_0.4_,_0.3_) optimize mate allocation by maximizing the index combining breeding value (BV) and Mendelian sampling variance under various constraints (0.6, 0.4, and 0.3) on the standardized haplotype similarity of parents. The results maximizing the BV are presented in Figure S7 in File S3. Results are reported for 100 simulation runs.

To better understand the effects of similarity matrices on genetic gain and variance, we quantified favorable QTL allele loss, SNP allele loss, the frequency of favorable alleles, and the number of selected sires (Figures 5 and S8). The OMA schemes involving the standardized similarity matrix preserved significantly more favorable QTL alleles than TS schemes (Figure 5A), preserving 5 – 19% more QTL alleles than TS on the index (Figure 5A). This enabled the OMA schemes to increase the frequency of these favorable alleles beyond those of their TS counterparts (Figure 5B). In addition, the OMA schemes retained significantly more SNPs than the TS schemes (Figure 5C), reflecting the selection and drift effect near QTL and neutral loci. Specifically, OMA schemes retained 3 – 19% more SNPs alleles than TS in the index (Figure 8C). The TS on BV resulted in a 1% higher rate of inbreeding than the TS on the index, which had an 8 – 26% higher rate of inbreeding than OMA schemes (Figure 5D). Although less pronounced, the unstandardized matrix similarly affected these quantities (Figures S9 and S10 in File S3).

**Figure 5.**
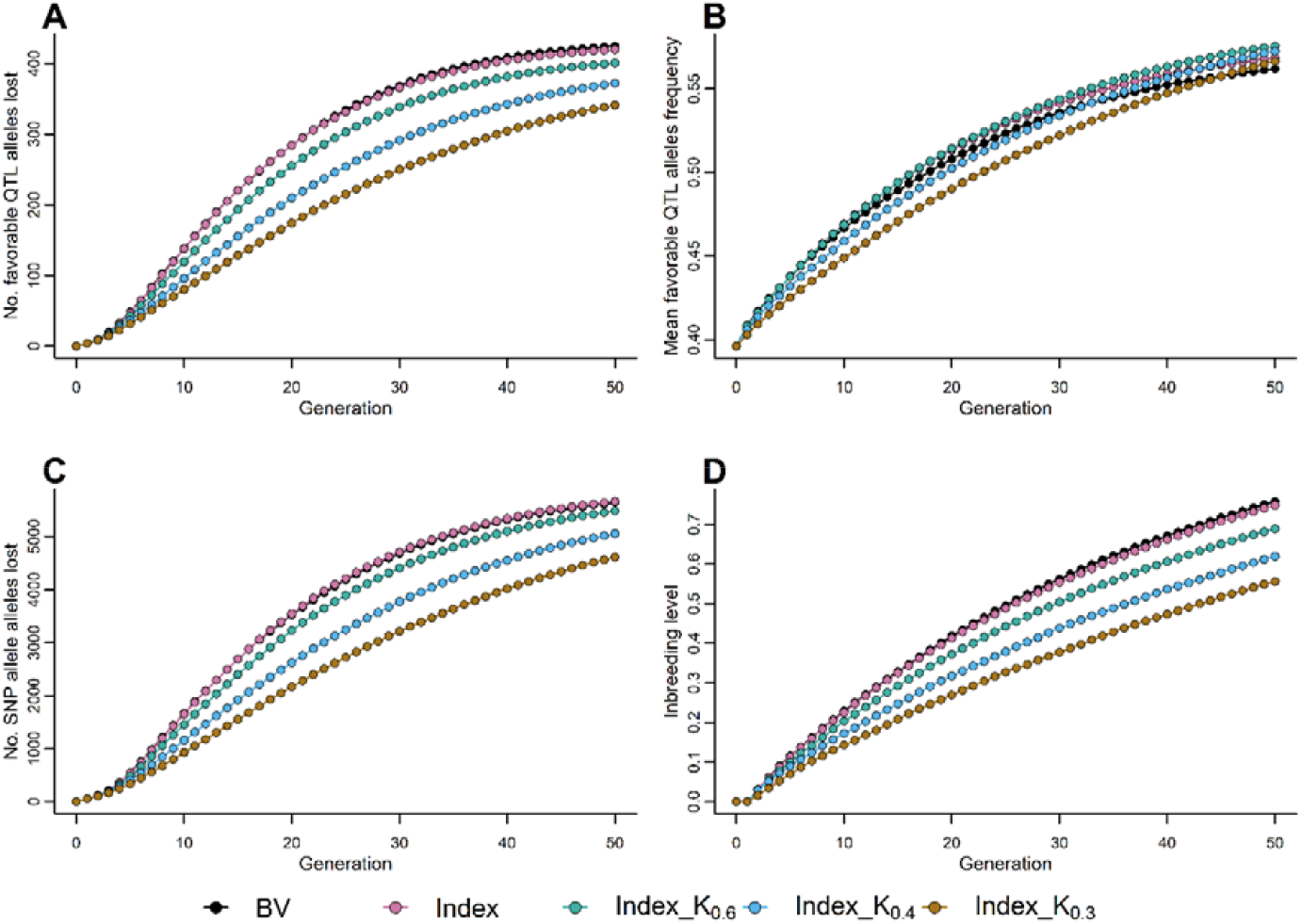
Effect of standardized similarity matrix (K) on favorable QTL alleles lost (A), mean favorable QTL allele frequency (B), SNPs lost (C), and expected inbreeding rate (D). The selection schemes Index_S_0.6_(_0.4_,_0.3_) optimize mate allocation by maximizing the index combining breeding value (BV) and Mendelian sampling variance under various constraints (0.6, 0.4, and 0.3) on the standardized haplotype similarity of parents. The results maximizing the BV are presented in Figure S8 in File S3. Results are reported for 100 simulation runs.

As the constraint on the average haplotype similarity of the parents increased, the number of selected sires for mating increased (Table S3 in File S3). Consequently, the diversifying effects on haplotypes in a group of selected parents also increased, preserving more favorable haplotypes. On average, OMA using the standardized similarity matrix selected significantly more sires than the non-standardized version.

## Discussion

The novel similarity matrix described in this paper fills a crucial methodological gap by providing a diversifying effect on haplotypes in a group of selected parents in cases where the selection criterion is either the breeding values or an index that combines the expected breeding values of offspring with their MSVs. Moreover, standardized similarities across the entire range of possible values appeared for several traits in real data, further underpinning the practical relevance of the approach. Notably, the derived similarity measure has a straightforward genetic interpretation—high similarity between parents arises from many shared chromosome segments with markers of large additive effects in the same linkage phase. When parents have identical haplotypes, the result would be a similarity that translates into a correlation of one. Possible alternatives for comparing Mendelian sampling values could be the Kullback–Leibler and Jensen–Shannon divergence methods, as known from information theory (Kullback and Leibler 1951; Nielsen 2019). However, their applicability to breeding was not investigated in this study.

One of the perks of our approach is the faster prediction of MSVs. We used marker effects with parent-specific signs and expected within-family LD matrix, unlike Bonk *et al*. (2016) and others (Santos *et al*. 2019; De Abreu Santos *et al*. 2020). Our approach took significantly less time to compute MSVs, and we expect the difference to be even greater when more parents and markers are involved. However, this computational advantage would be lost if parent-specific recombination rates or marker distances became available.

The similarity matrices’ derivations are based on the availability of phased genotypes, a genetic map, and estimated marker effects. The accuracy of these data is expected to affect the entries of a similarity matrix, but the magnitude of this effect is still unclear. The similarity matrices are trait-specific and require no weighting in a selection involving only a single trait. However, index weights allow all trait-specific similarity matrices to be summed when selecting multiple traits.

Visualization is one of the many applications of similarity matrices among parents. Using the Holstein-Friesian dataset, we showed how these similarity matrices provide a quick overview of the similarity of Mendelian sampling values within and across families. The size of these similarities depends on the linkage and heterozygosity in parents and the trait genetic architecture (Figures 1–2 and S4–5 in File S3). Standardized similarities also have an interpretation as the proportion of shared MSV. The observed standardized pairwise similarities for all traits in the empirical data of the present study ranged almost across the entire possible parameter space (Figure 2). However, we expect the species and populations of plants and animals to affect these values. Their influence on the similarities of Mendelian sampling values should be investigated in future studies.

To demonstrate the application of the similarity matrices in hedging haplotype diversity, we compared several selection schemes: combinations of BV or index with TS or OMA rule. The OMA schemes maximized the breeding value or index under a given constraint on parents’ average haplotype similarity, assuming known marker effects and sex-averaged genetic map. The longer-term genetic gain and greater variance preserved by TS on the index over BV (Figure 3) were expected due to the index’s better preservation of favorable QTL alleles (Figure S5) and consistent with previous simulation studies using similar indices (Müller *et al*. 2018; Allier *et al*. 2019; Musa and Reinsch 2022). The greater long-term genetic gain and variance realized by OMA schemes than their TS counterparts, at the expense of lower short-term genetic gain (Figure 3 and Figure S6 in File S3), was also expected, as it is the typical trend with long-term genomic selection strategies (Daetwyler *et al*. 2015; Liu *et al*. 2015; Müller *et al*. 2018; Allier *et al*. 2019). This is because long-term genomic selection strategies sacrifice selecting the most performant candidate parents to preserve genetic diversity and achieve long-term gains. What we found intriguing, however, was the lower short-term genetic variance in OMA schemes (Figure 3B and S6B in File S3), which can be attributed to the similarity measure.

The similarity measure is designed to help prevent the selection of similar parents with high MSV potential to avoid unintended loss of favorable haplotypes when selection is based on the index or breeding values. Constraining the haplotype similarity of parents may therefore sacrifice the selection of parents with large MSVs caused by favorable haplotypes in coupling with large effects on the trait(s) of interest in favor of parents with lower MSVs resulting from different chromosomal segments. The former could recombine in later generations to produce high genetic variability. In contrast, despite having parents with more diverse haplotypes, the latter could produce lower genetic variability in initial generations, as observed in Figures 3B and S6B in File S3.

On the other hand, constraining the standardized similarity measure emphasizes the similarity between parental haplotypes rather than MSVs, allowing a greater range of favorable haplotypes to remain in play over successive generations (Figures 5A and S8A in File S3). As a result, up to 7% and 1630% more long-term genetic gain and variance were realized than index selection (Figures 4 and S7 in File S3), in contrast to constraining the unstandardized similarity measure, which resulted in only up to 2% and 312% more genetic gain and variance. The standardized similarity matrix also results in a less conspicuous lower short-term genetic variance (Figures 4B and S7B in File S3) than observed with the unstandardized version.

The OCS (Meuwissen and Sonesson 1998; Sonesson *et al*. 2012; Woolliams *et al*. 2015) and OMA strategies aim to optimize parental contributions to the next generation by maximizing genetic gain under a given constraint on the average similarity between parents. They, however, differ in two key ways. Firstly, OCS is only applicable to individuals, whereas OMA can also be applied to parent pairs (see Section “Similarities between zygotes of two pairs of parents”). Second, and more importantly, the similarity measure for OCS is a GRM based on parental genotypes, whereas the similarity measure for OMA is based on Mendelian segregation patterns. Thus, the similarity measures account for various facets of diversity and serve distinctly different functions.

The OCS has been found to either maintain allele frequencies close to the original population or shift them toward 0.5, depending on the GRM used (Gómez-Romano *et al*. 2016; Meuwissen *et al*. 2020). Because genetic improvement is driven by increasing the frequency of favorable alleles, OCS may not efficiently maximize short and mid-term genetic gain and could accumulate deleterious alleles in the genome, thereby reducing population fitness (de Cara *et al*. 2013). Consequently, a similarity matrix indicating the degree to which parents share heterozygous QTL segments used in OMA schemes may be a more appropriate similarity measure when selection aims to maximize genetic gain while preserving genetic diversity.

To test this hypothesis, we compared our results with those of optimizations involving GRM, henceforth referred to as OCS schemes. We developed additional selection schemes with ***Q***_***t***_ as GRM and constraints of 1% and 0.5% on the average rate of inbreeding (see Eq. 18). GRMs were derived according to Method 1 of VanRaden (2008) and a constraint was placed such that a minimum of 5 males would be selected for mating. As with OMA schemes, only males were optimized in OCS schemes, while females were selected based on either TS on BV or index. We found that the OCS schemes generally resulted in greater long-term genetic gain and preserved more genetic variability than the OMA schemes at the expense of short-term genetic gain (Figure S11A and B in File S3). The greater variability preserved could result from better control of the inbreeding rate, which led to the selection of a mean of 35 – 72 sires across OCS schemes, whereas OMA schemes selected 8 – 20 sires (Table S4 in File S3). To determine the effect of combining both matrices on genetic gain and variance, we added a 0.5% constraint on the average inbreeding rate to OMA schemes, i.e., 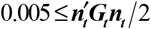, where ***G*** is the GRM. This combination improved long-term genetic gain and preserved more variability relative to the OMA schemes. It also preserved equal or greater long-term genetic variance relative to OCS schemes (Figure S11C and D in File S3), selecting a mean of 71 – 72 sires across generations (Table S4 in File S3).

We further increased the minimum number of selected sires to 25 and discovered that OCS selected a mean of 35 – 71 sires, OMA schemes selected 25 – 33 sires, and the combination of both matrices selected 70 – 71 sires across generations (Table S4 in File S3). Here genetic gain and variance were comparable in the long term, and where differences existed, OCS had the advantage over OMA schemes (Figure S12 in File S3). The reverse was true when the minimum number of sires to be selected further increased to a minimum of 50 sires (Figure S13 in File S3). In every scenario, combining both matrices enhanced the preservation of genetic variability in the long term, indicating that GRM and our similarity matrices cover different aspects of diversity and, more importantly, complement each other.

It should be noted that OMA can be improved by tweaking constraints, non-random mating, and optimizing the selection of males and females as opposed to only males (as was the case in the present study). Thus, extensive studies are required in scenarios that vary these parameters, genetic architecture, and selection for multiple traits to compare OMA with other strategies and determine its strengths and weaknesses. Further study is required on other technical aspects of OMA, such as recommending constraints for various populations. The recommended constraints in OCS are 0.5 – 1% inbreeding rates for livestock populations (FAO 2015; Meuwissen and Oldenbroek 2017). However, recommending constraints for OMA is challenging because the similarity matrices’ values are highly dependent on factors such as genetic architecture and the number of traits under selection, which have no bearing on OCS since GRM is based on genome-wide similarities.

Another technical aspect involves dealing with an indefinite similarity matrix when it is encountered. Apart from floating point inaccuracies, one reason for indefiniteness could be rank deficiencies in the family-specific marker effects vector ***m*** _*i*_. Recall that homozygous loci in ***m*** _*i*_ vectors are set to zero and that heterozygous markers retain the marker effects at respective loci, but the sign varies based on the linkage phase. This implies that heterozygous loci with zero marker effects are also zero. Thus, the ***m***_*i*_ vectors (depending on the data) simulate a situation where the number of parents is far greater than the number of informative markers. This phenomenon is amplified because similarities are calculated chromosome-by-chromosome, which holds the number of parents constant while varying the number of informative markers (often smaller than the number of parents). See Table S5 in File S4, which shows the properties of the similarity matrix for each chromosome and genome for a randomly chosen simulation run. Additional contributing factors include heterozygosity of parents, genetic trait architecture, marker effects estimation method, population type, and number of traits under selection.

In conclusion, we derived trait-specific similarities between parents reflecting the shared heterozygosity of their markers with large effects and similar linkage phases. The similarity of a parent to itself is equal to its MSV. When combined in a matrix, these similarities provide the opportunity to balance haplotype diversity and genetic gain in selection decisions. Combining the similarity matrix and breeding value or index preserved genetic variability in a simulated breeding program more effectively than selection based on breeding value and index alone without compromising genetic gain. The combination of similarity matrix and index better preserved genetic variance and resulted in higher genetic gain than the combination with breeding value. In addition, the combination of the similarity matrix and the genomic relationship matrix preserved even more genetic variance because they complemented each other. This study focused on applying the similarity matrices in optimizing mate allocation when genotypic data from both male and female parents are available. However, they can also be used to optimize mating decisions in scenarios where genotypic data are only available for one sex. Another potential application of gametic similarity matrices is plant breeding, where gametes from parents are converted into double-haploid lines that form the next breeding generation.

## Data availability

The datasets supporting the conclusions of this article are included in the article and its supplementary material. The pedigree information, genotypic data, and physical map of the empirical dataset can be found here: DOI: <u>10.3389/fgene.2018.00186</u> under the Supplementary Material section.

## Acknowledgments

The empirical data provided by Nina Melzer and Dörte Wittenburg is gratefully acknowledged. Dörte Wittenburg’s suggestions for improving the manuscript are greatly appreciated.

## Supplemental data

Supplemental material – zip file

## Appendices

### Appendix 1: Illustration of equivalence

Eqs. (2) and (5) are equivalent. To demonstrate their equivalence, consider the following case of two biallelic marker loci on a single chromosome with haplotypes in coupling and repulsion phases.

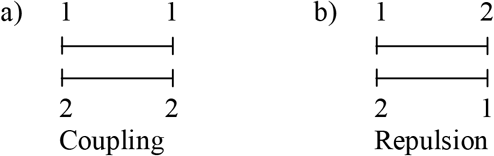

According to Bonk *et al*. (2016), for any two-marker interval, we have the following covariance matrices:

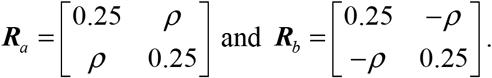

Meanwhile, the MSVs are as follows:

a. 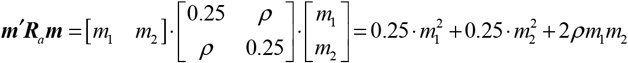
b. 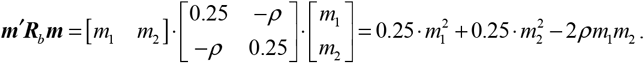

After setting up ***m***_*a*_ and ***m***_*b*_, the same results can be obtained by setting up a population covariance matrix

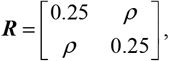

and applying Eq. (5):

a. 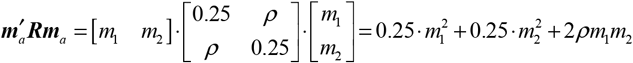
b. 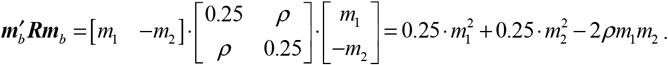.

The equivalence of the two representations of MSV generalizes to any number of markers (see Appendix 2 for proof of equivalence).

### Appendix 2: Proof of equivalence

In Eq. (2) 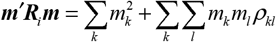 each *ρ*_*kl*_ is positive if the marker alleles to which the effects *m*_*k*_ and *m*_*l*_ have been assigned are on the same chromosome (in coupling). Otherwise, *ρ*_*kl*_ is negative. As the marker effects do not depend on the linkage phase, each product *m*_*k*_ *m*_*l*_ *ρ*_*kl*_ changes its sign according to the linkage phase of the respective markers.

In Eq. (5) 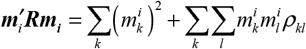 the covariance *ρ*_*kl*_ is always positive, but the signs of the marker effects depend on the linkage phase. In coupling, they are as displayed in Eq. (2), yet in the linkage phase, they are either (*m*_*k*_, −*m*_*l*_) or (−*m*_*k*_, *m*_*l*_) depending on the alleles on the first haplotype. In effect, this causes the sign of each product 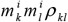 to change according to the linkage phase as before and 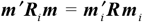. Thus, it is quicker to set up ***m***_*i*_ with *M* elements on a chromosome than 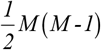 off-diagonal elements of a half-stored ***R*** -matrix.

### Appendix 3: Monoecious species and order of parents in dioecious organisms

In monoecious plants, parents *i* and *j* may swap their function as male (pollen donor) and female parents when producing offspring. The MSVs of the resulting zygotes then remain the same, assuming equal recombination rates in male and female parents. With sex-specific covariance matrices ***R***_*m*_ and ***R*** _*f*_ reflecting maps of males and females, the MSVs for parents *ij* and *ji* are

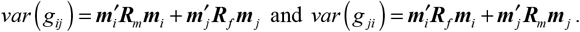

For two pairs of parents *ij* and *ji* there are, in principle, four different conditional covariances, depending on which individual acts as pollen donor within each pair:

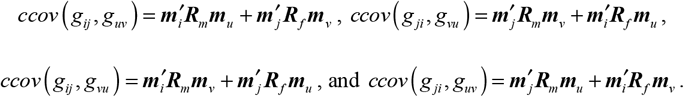

If all four kinds of parentage have equal probabilities and 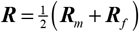, then the average of these four conditional covariances can be simplified as

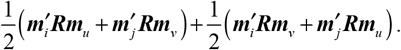

Then, by taking absolute values in a chromosome-wise manner as described above, this average conditional covariance defines the similarity 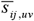 between parents *ij* and *uv* as

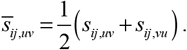

This could also be used as an alternative similarity measure in mammals and dioecious plants if the similarity is to be independent of the order of parents—or, equivalently, independent of the parental origin of the haplotypes in the offspring.

