## Supplementary file 1 for "A similarity matrix for preserving haplotype diversity among parents in genomic selection"

**Derivation of Mendelian sampling variance, covariance, and similarity from a probability distribution**

Mendelian sampling is the independent segregation of alleles of various genes into gametes, provided that the genes belong to different linkage groups. Genes on the same chromosome belong to the same linkage group. The tendency of the alleles of genes on the same chromosome to recombine into gametes can be measured from a mapping function (e.g., Haldane 1919). The segregation patterns of alleles into gametes can be obtained straightforwardly using a probability tree representing a sequence of events.

### A. Gametes

To demonstrate, we assume three biallelic markers on a single chromosome and two vectors of population additive marker effects for trait 1
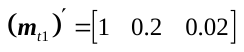
 and trait 2
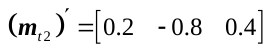
. Five potential parents with phased haplotypes—coded as
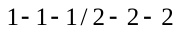
 for the first,
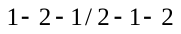
 for the second,
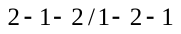
 for the third,
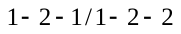
 for the fourth, and
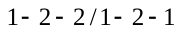
 for the fifth parents—are assumed. Finally, we assume a vector of genetic distances (Haldane Morgan) between markers
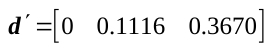
, which translates to recombination rates of
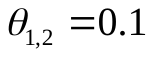
 between marker loci 1 and 2 and
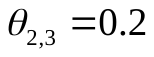
 between loci 2 and 3.

Figure S1 represents all segregation patterns of alleles into gametes, probabilities of occurrence, and the genetic values for trait 1 of the first and second parents. From the probability tree, it is clear that gametes from two parents have a similar probability of occurrence, but they usually have different genetic values, depending on the parent’s genetic makeup. The genetic values obtained from different segregation patterns of gametes for trait 1 of the first and second parents (
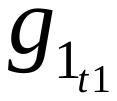
 and
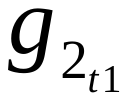
) are derived from parent-specific vectors (explained in Eq. (5)), and they represent the value transmitted to the offspring (also see Table 1).

#### A.1 Mendelian sampling variance of gametes

From the probability distribution of segregation patterns, the Mendelian sampling variances (MSVs) of genetic values for the first (
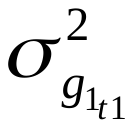
) and second (
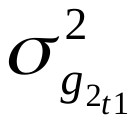
) parents for trait 1 (
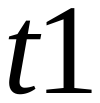
) are
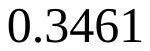
 and
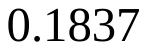
, respectively (see Table 1). Alternatively, and more conveniently, the MSV for parent
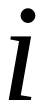
 can be obtained using the equivalent matrix formula
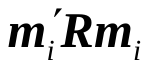
 (Eq. 5)

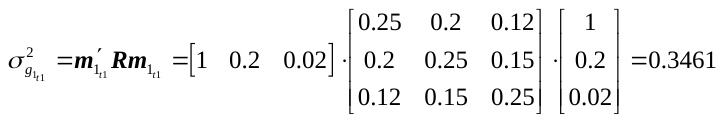

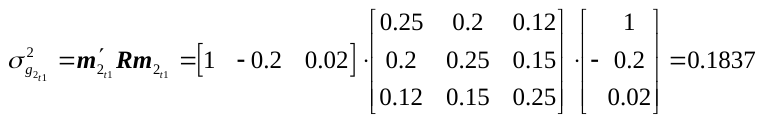
,

where
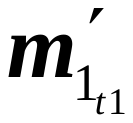
 and
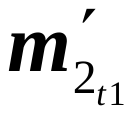
 are row vectors of parent-specific additive marker effects for trait
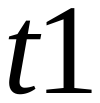
 of the first and second parents, respectively. MSV does not depend on the haplotype order chosen when setting up the parent-specific vector
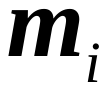
.

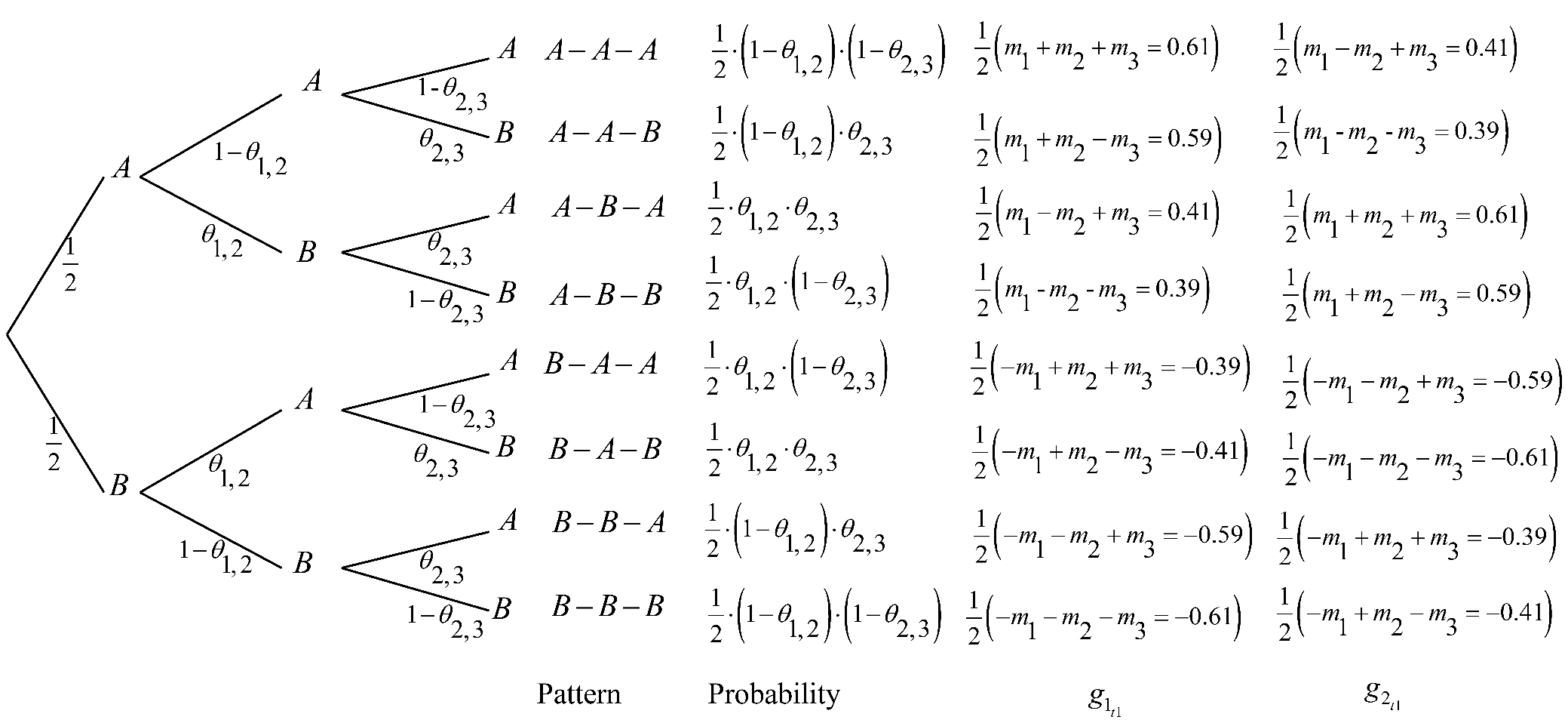

**Figure S1** A probability tree showing the segregation patterns (sequence of alleles from the first (
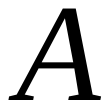
) and second (
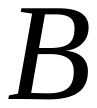
) parental haplotypes), probabilities of occurrence, and the genetic values for trait
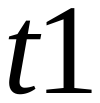
 of the first (
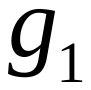
) and second (
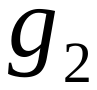
) parents.

#### A.2 Mendelian sampling covariance for multiple traits of the same parent

Figure S2 shows the segregation patterns, probabilities of occurrence, and genetic values of gametes produced by the first parent for multiple traits. The Mendelian sampling covariance (MSC) between gametes for multiple traits
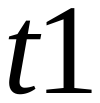
 and
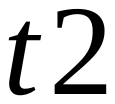
 of the first parent can be derived from the following probability distribution:

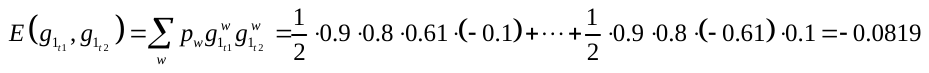

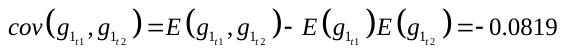
.

Here
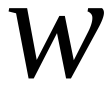
 is the index of the segregation pattern,
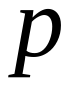
 is the probability of segregation pattern,
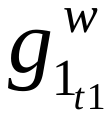
 and
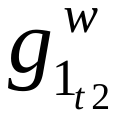
 are the additive value of gamete transmitted from the first parent for traits
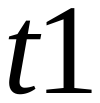
 and
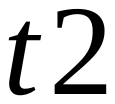
.

Alternatively, the same results can be obtained using
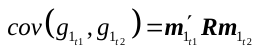
, where
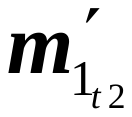
 is the parent-specific vector of marker effects for trait
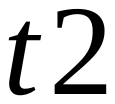
 of the first parent. Applying the formula
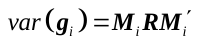
 (Eq. 6), we obtain

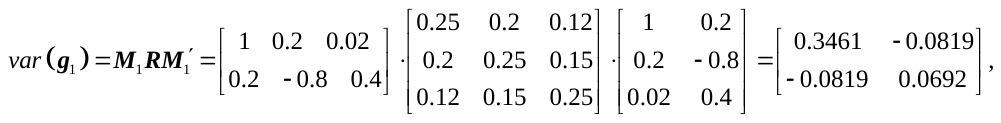
where the rows of matrix
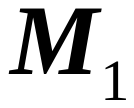
 are parent-specific vectors for each trait (
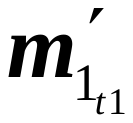
 and
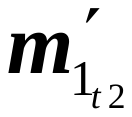
) of the first parent. The diagonal elements of
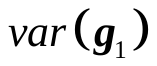
 are the MSVs of each trait, whereas the off-diagonal elements are the MSCs. Like MSV, the MSC between multiple traits does not depend on the haplotype order chosen when setting up the parent-specific vector
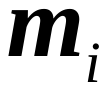
.

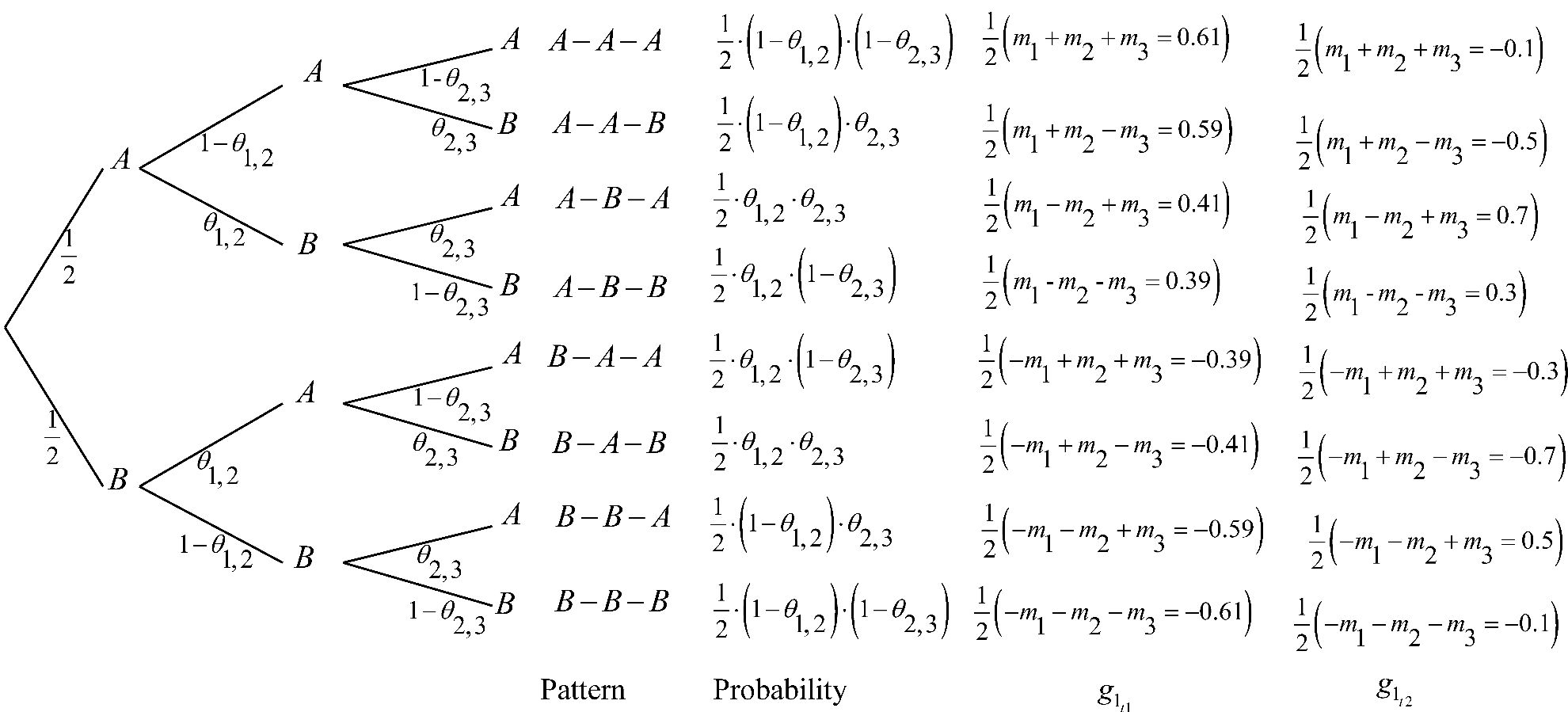

**Figure S2**: A probability tree showing the segregation patterns (sequence of alleles from first (
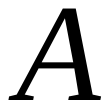
) and second (
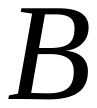
) parental haplotypes), probabilities of occurrence, and the genetic values for multiple traits (
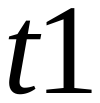
 and
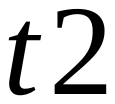
) of the first parent (
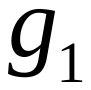
).

#### A.3 Conditional covariance of Mendelian sampling values of gametes from different parents

The Mendelian sampling processes in two parents are random and independent. Enumerating the covariance between two parents from a bivariate distribution of gametes with probabilities of all combinations for possible gametes would yield a covariance of zero (Table S1). Table 1 and Figure S1 show that the probability distribution gives the frequencies of all possible segregation patterns of gametes (
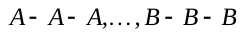
) within a family. Assuming identical probabilities of the within-family distribution of segregation patterns of gametes from two parents—which usually have different genetic values depending on their genetic makeup—we can obtain a non-zero covariance between two parents. However, this is conditional on the chosen order of haplotypes in the parent-specific vector
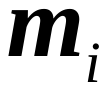
.

From a probability distribution, the derivation of conditional covariance between Mendelian sampling values for trait
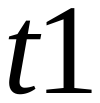
 of the first and second parents is
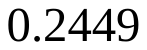
 (Table 1). The same result can be obtained using
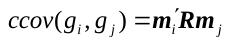
:

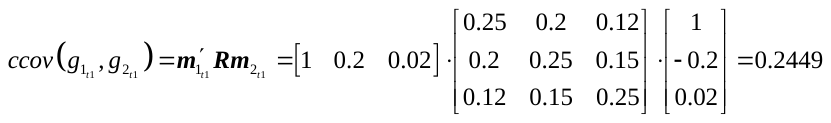
.

The vectors of parent-specific additive marker effects for trait
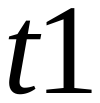
 of the first and second parents have been given above. For the third, fourth, and fifth parents, the vectors are
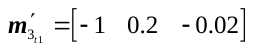
,
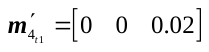
, and
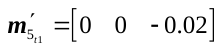
, respectively. From this, a comprehensive conditional covariance matrix for all five parents for a single trait
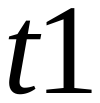
 can be produced:

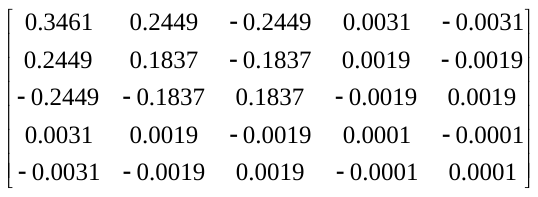
*.*

The MSVs of all parents constitute the diagonal elements of the matrix above, whereas the off-diagonal elements represent the conditional covariance between the Mendelian sampling values of the parents. As seen in the matrix, the covariance between parents depends on the order of haplotypes. The conditional covariance between the first and second parents is 0.2449, while it is -0.2449 between the first and third parents, indicating that the second and third parents share similar heterozygosity but entirely swapped haplotype order.

Therefore, the similarity between parents
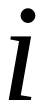
 and
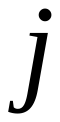
 should be calculated using the formula
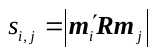
 (Eq. 9) to obtain an unequivocal value of similarity independent of the order of haplotypes. Applying Eq. (9) and assembling all pairwise similarities into a matrix, we obtain the following similarity matrix:

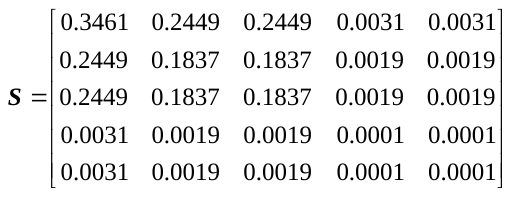
.

Standardizing matrix
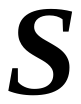
, according to Eq. (17), yields:

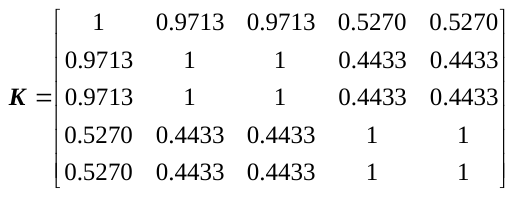
.

Matrices
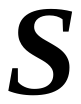
 and
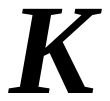
 show that parents with common heterozygosity of markers with large effects (first and second parents) appear to be more similar than those with small effects (first and fifth parents).

| Table S1: Bivariate probability distribution of segregation patterns of gametes in parents 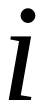 and 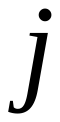 | | | | | | | | |
| --- | --- | --- | --- | --- | --- | --- | --- | --- |
| Parent 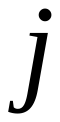  Parent 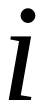 | 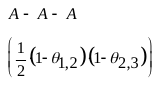 | 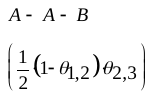 | 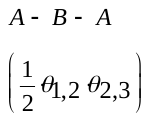 | 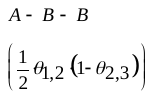 | 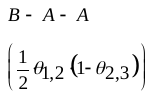 | 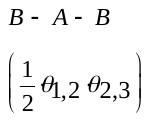 | 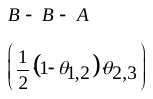 | 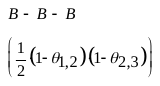 |
| 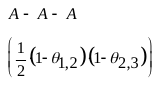 | 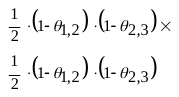 |  |  |  |  |  |  |  |
| 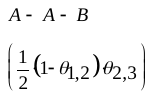 | 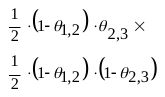 | 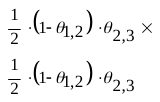 |  |  |  |  |  |  |
| 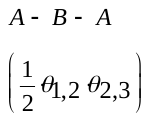 | 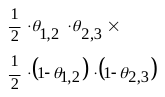 | 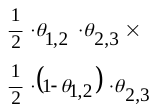 | 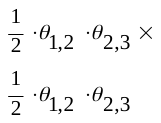 |  |  |  |  |  |
| 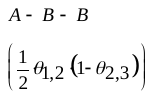 | 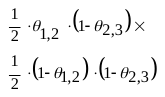 | 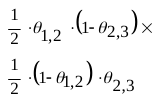 | 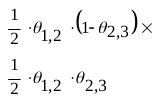 |  |  |  |  |  |
| represents alleles on the first haplotype; represents alleles on the second haplotype. Only the lower triangle is shown. | | | | | | | | |

### B. Zygotes

#### B.1 Similarity between Mendelian sampling values of zygotes

The makeup of haplotypes that zygotes inherit from male and female parents can also be characterized by the recombination patterns of parental haplotypes. and represent alleles from the first and second haplotypes of a parent, respectively. Then, a zygote with the segregation pattern combines a recombinant haplotype from the first parent and a non-recombinant one from the second. Two pairs of parents can produce two populations of zygotes that match into pairs by their recombination pattern, depending on the haplotype orders chosen for the parents. The zygotes of each pair share the same probability of occurrence, but their genetic values may differ depending on their parents’ genetic makeup. From the probability distribution of segregation patterns, we can compute the conditional covariance among the genetic values of zygotes produced by their parents.

Consider two linked biallelic marker loci in the zygotes produced by four parents, , , , and , with haplotypes , , , and , respectively. Assuming a vector of population additive marker effects and a recombination rate of (0.1116 Haldane Morgan), the matching common segregation patterns, probabilities of occurrence, and genetic values of zygotes produced by parent pairs and are summarized in Table S2 and Figure S3.

| **Table S2 Segregation patterns, probabilities of occurrence, marker haplotypes, and genetic values of zygotes produced by two pairs of parents (** **and** **) with matching recombination patterns** | | | | | | | |
| --- | --- | --- | --- | --- | --- | --- | --- |
| **Segregation pattern/**  **Zygotes produced** | | **Probability** | **Marker haplotypes of zygotes** | |  | **Genetic value of zygotes** | |
|  |  |  |  |  |  | **Parents** | **Parents** |
| A represents the alleles on the first haplotype, and B represents alleles on the second haplotype of a zygote parent. Parents , , , and have ordered haplotypes , , , and , respectively. A recombination rate of and a vector of population additive marker effects are assumed. Pattern , for example, has alleles in parent pair and in parent pair . Note that genetic value represents the value transmitted to the offspring. | | | | | | | |

The variance of the genetic values of the zygotes produced by parent pair can be derived from the probability distribution of segregation patterns as follows:

.

Following the same procedure, the variance of the genetic values from zygotes produced by parent pair is .

Similar to the gametic case, the similarity between the zygotes of parent pairs and can be derived from the probability distribution of joint segregation patterns (probabilities ) as follows:

.

The same results for MSV can be obtained using matrix expression (Eq. 11),

,

and for conditional covariance using (Eq. 15),

.

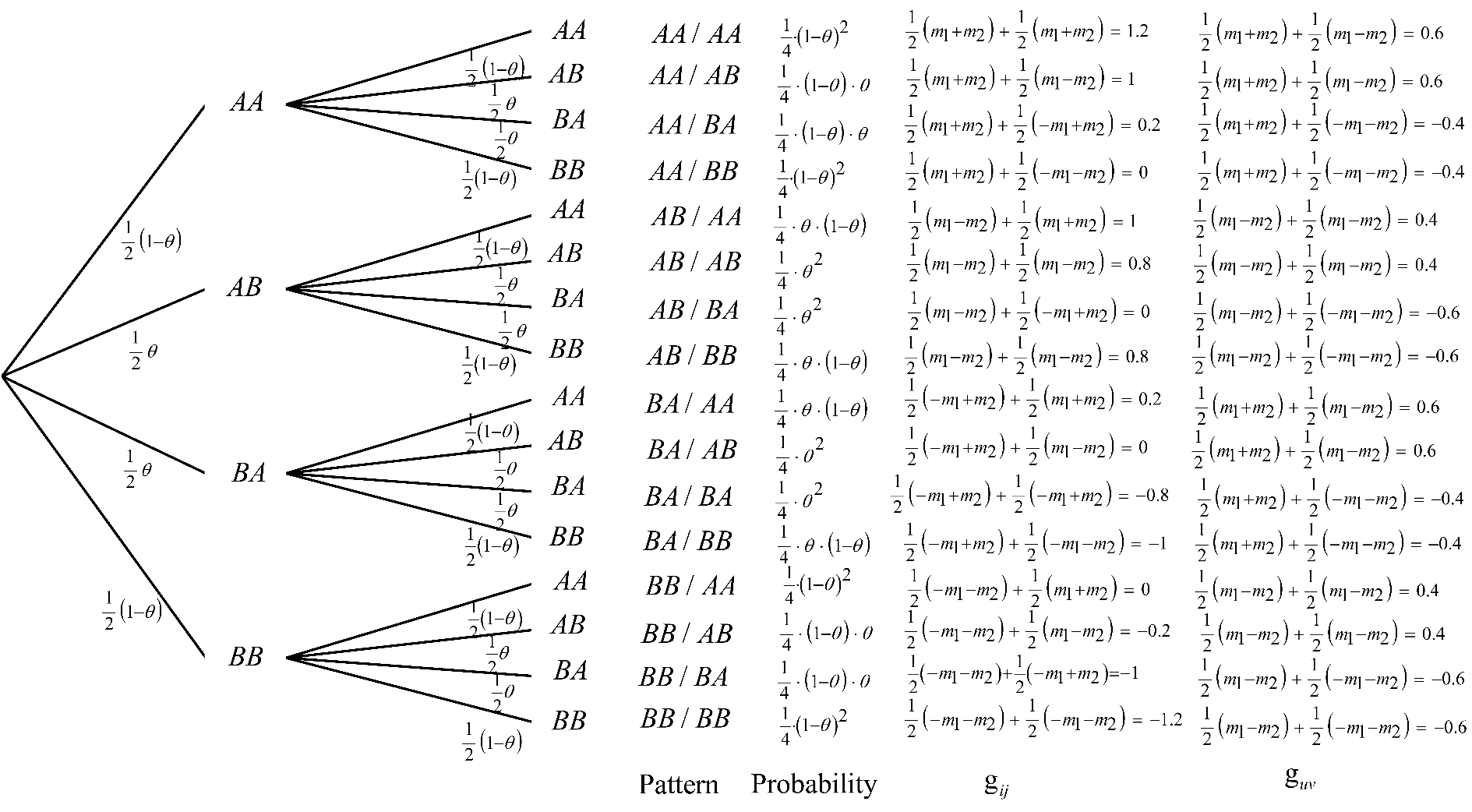

**Figure S3**: A probability tree showing the segregation patterns (sequence of alleles from first () and second () parental haplotypes), probabilities of occurrence, and the genetic values of zygotes from parent-pairs and .
