## Supplementary file 3 for "A similarity matrix for preserving haplotype diversity among parents in genomic selection"

**TABLES**

| **Table S3: Summary Statistics for the number of selected males by various selection schemes** | | | |
| --- | --- | --- | --- |
| **Scheme*** | **Minimum** | **Mean** | **Maximum** |
| Optimum mate allocation schemes involving similarity matrix | | | |
| BV_S_0.6_ | 5 | 6.1 | 17 |
| BV_S_0.4_ | 5 | 8.2 | 20 |
| BV_S_0.3_ | 5 | 9 | 21 |
| Index_S_0.6_ | 5 | 7 | 18 |
| Index_S_0.4_ | 5 | 9.5 | 22 |
| Index_S_0.3_ | 5 | 10.5 | 22 |
| Optimum mate allocation schemes involving standardized similarity matrix | | | |
| BV_K_0.6_ | 5 | 8.3 | 20 |
| BV_K_0.4_ | 5 | 14.5 | 30 |
| BV_K_0.3_ | 7 | 18.6 | 31 |
| Index_K_0.6_ | 5 | 9.6 | 19 |
| Index_K_0.4_ | 7 | 16.1 | 29 |
| Index_K_0.3_ | 9 | 20.4 | 35 |
| *Truncation selection on BV and index always selected five males. BV = breeding values; Index = combination of BV and Mendelian sampling variance; S = similarity matrix; K = standardized similarity matrix. The indices 0.6, 0.4, and 0.3 represent the 60th, 40th, and 30th percentiles of haplotype similarities of the base population and correspond to 0.18, 0.15, and 0.14 constraints imposed on S and 0.40, 0.35, and 0.33 constraints imposed on K, respectively. | | | |

| **Table S4: Summary Statistics for the number of selected males by various selection schemes** | | | | | | | | | | | |
| --- | --- | --- | --- | --- | --- | --- | --- | --- | --- | --- | --- |
| **Scheme*** | **Minimum** | | |  | **Mean** | | |  | **Maximum** | | |
|  | **5** | **25** | **50** |  | **5** | **25** | **50** |  | **5** | **25** | **50** |
| **Optimum mate allocation** | | | | | | | | | | | |
| BV_K_0.6_ | 5 | 25 | 50 |  | 8.3 | 25 | 50 |  | 20 | 29 | 51 |
| Index_K_0.6_ | 5 | 25 | 50 |  | 9.6 | 25.3 | 50 |  | 19 | 32 | 53 |
| BV_K_0.3_ | 7 | 26 | 50 |  | 18.6 | 32.1 | 55.4 |  | 31 | 43 | 64 |
| Index_K_0.3_ | 9 | 26 | 50 |  | 20.4 | 33.1 | 56 |  | 35 | 45 | 64 |
| **Optimum contribution selection** | | | | | | | | | | | |
| BV_G_0.01_ | 26 | 25 | 50 |  | 40.1 | 35.1 | 50 |  | 60 | 51 | 52 |
| Index_G_0.01_ | 27 | 25 | 50 |  | 40.1 | 35.4 | 50 |  | 59 | 49 | 53 |
| BV_G_0.005_ | 52 | 52 | 51 |  | 71.2 | 70.2 | 67.3 |  | 95 | 98 | 86 |
| Index_G_0.005_ | 54 | 54 | 51 |  | 72.1 | 70.9 | 67.7 |  | 92 | 96 | 89 |
| **Combining standardized similarity matrix and genomic relationship matrix** | | | | | | | | | | | |
| BV_K_0.6_G_0.005_ | 55 | 54 | 52 |  | 71.2 | 70 | 67.3 |  | 92 | 95 | 92 |
| Index_K_0.6_G_0.005_ | 54 | 53 | 54 |  | 72.2 | 71 | 67.8 |  | 92 | 92 | 88 |
| BV_K_0.3_G_0.005_ | 56 | 55 | 54 |  | 71.1 | 70.1 | 68.7 |  | 92 | 88 | 87 |
| Index_K_0.3_G_0.005_ | 52 | 56 | 53 |  | 71.5 | 70.4 | 69.1 |  | 92 | 88 | 85 |
| *Truncation selection on BV and index always selected five males. BV = breeding values; Index = combination of BV and Mendelian sampling variance; K = standardized similarity matrix; G = genomic relationship matrix. The indices 0.6, 0.4, and 0.3 of K represent the 60th, 40th, and 30th percentiles of haplotype similarities of the base population and correspond to 0.40, 0.35, and 0.33 constraints imposed, respectively. The indices 0.01 and 0.005 are the constraints imposed on G. | | | | | | | | | | | |

**FIGURES**

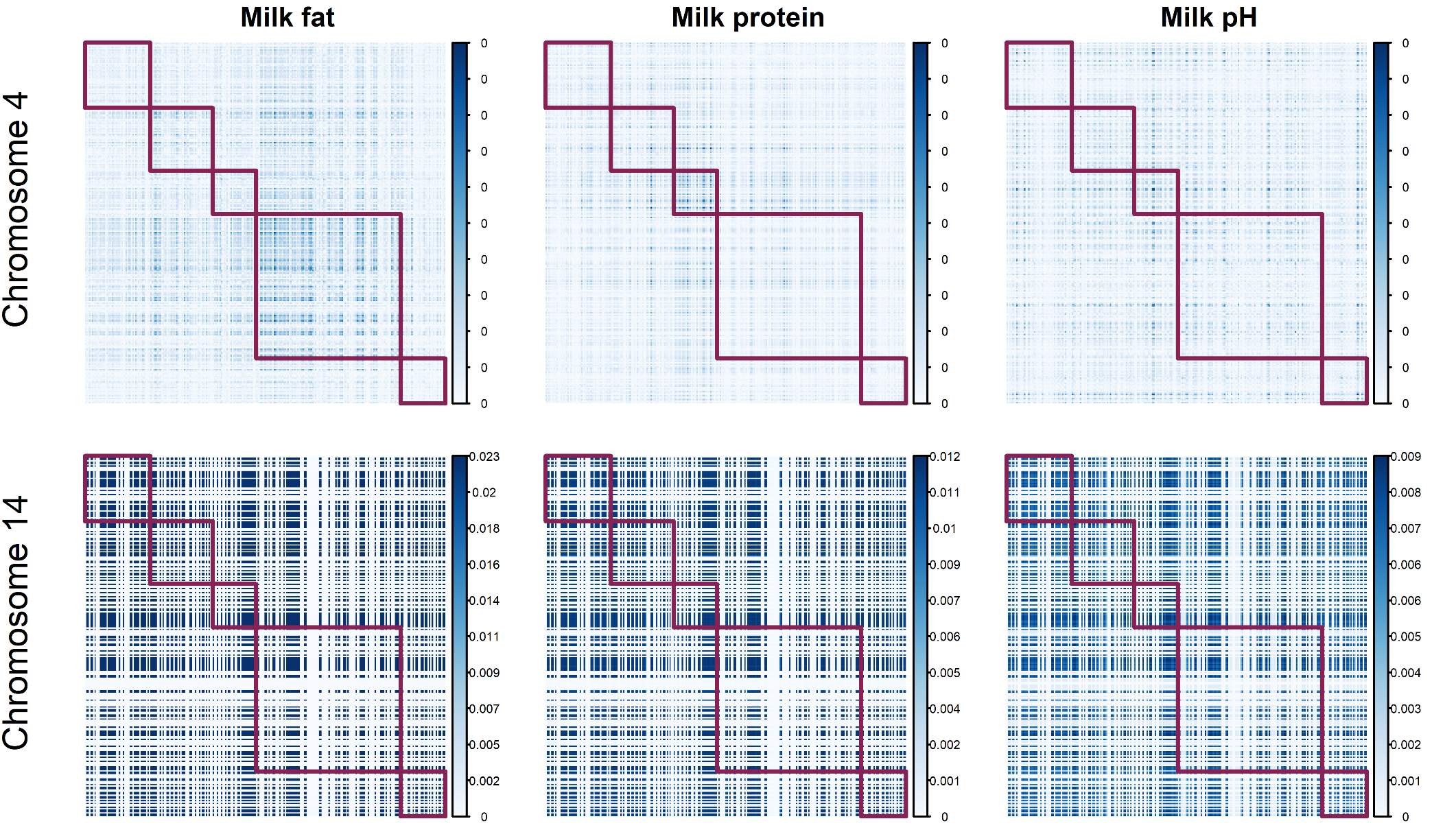

**Figure S4** Similarity matrices showing chromosomes 2 and 14 for three milk fat, protein, and pH. The red blocks demarcate each paternal half-sib family. Parents are arranged according to their pedigree.

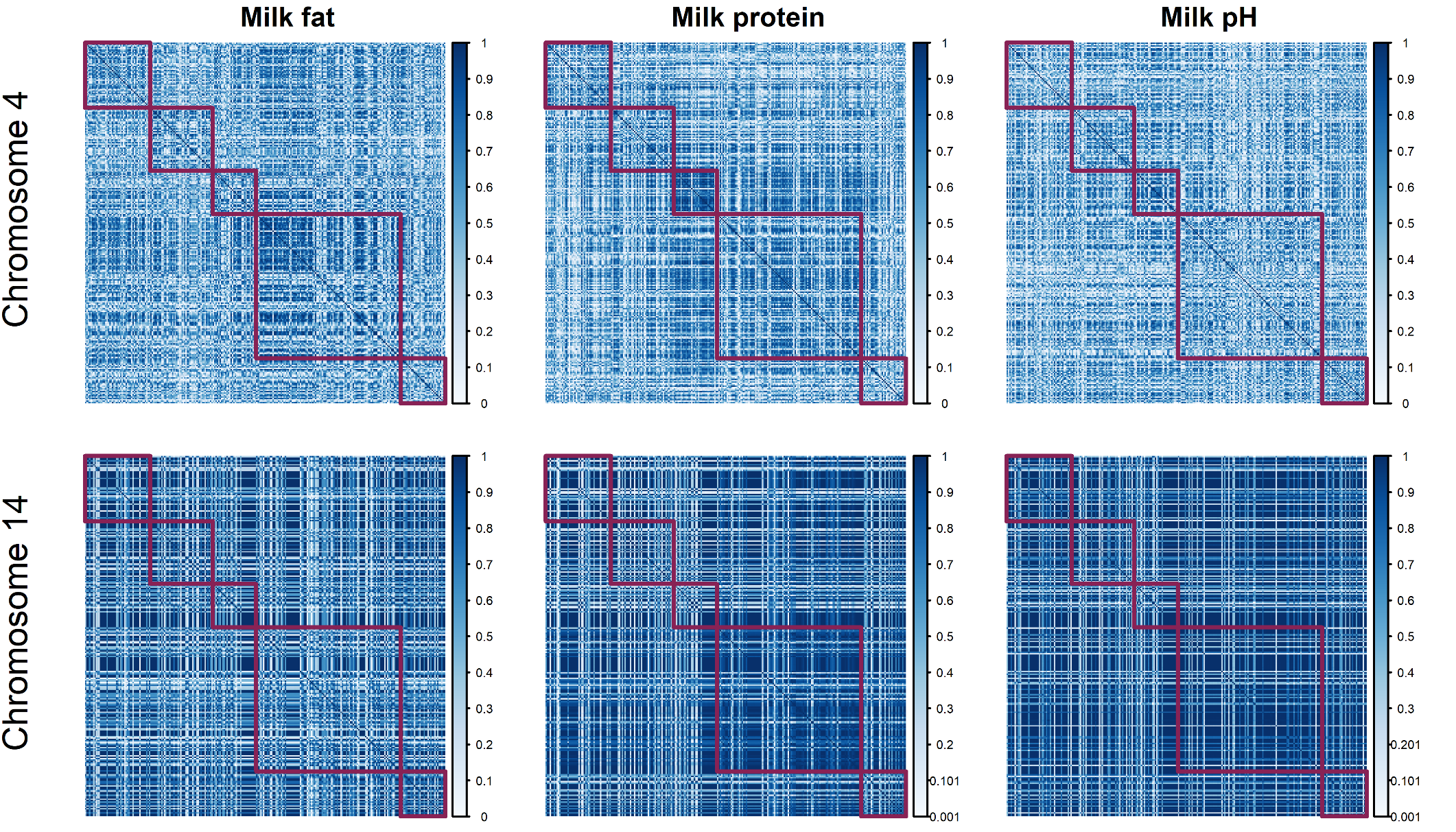

**Figure S5** Standardized similarity matrices showing chromosomes 2 and 14 for three milk fat, protein, and pH. The red blocks demarcate each paternal half-sib family. Parents are arranged according to their pedigree.

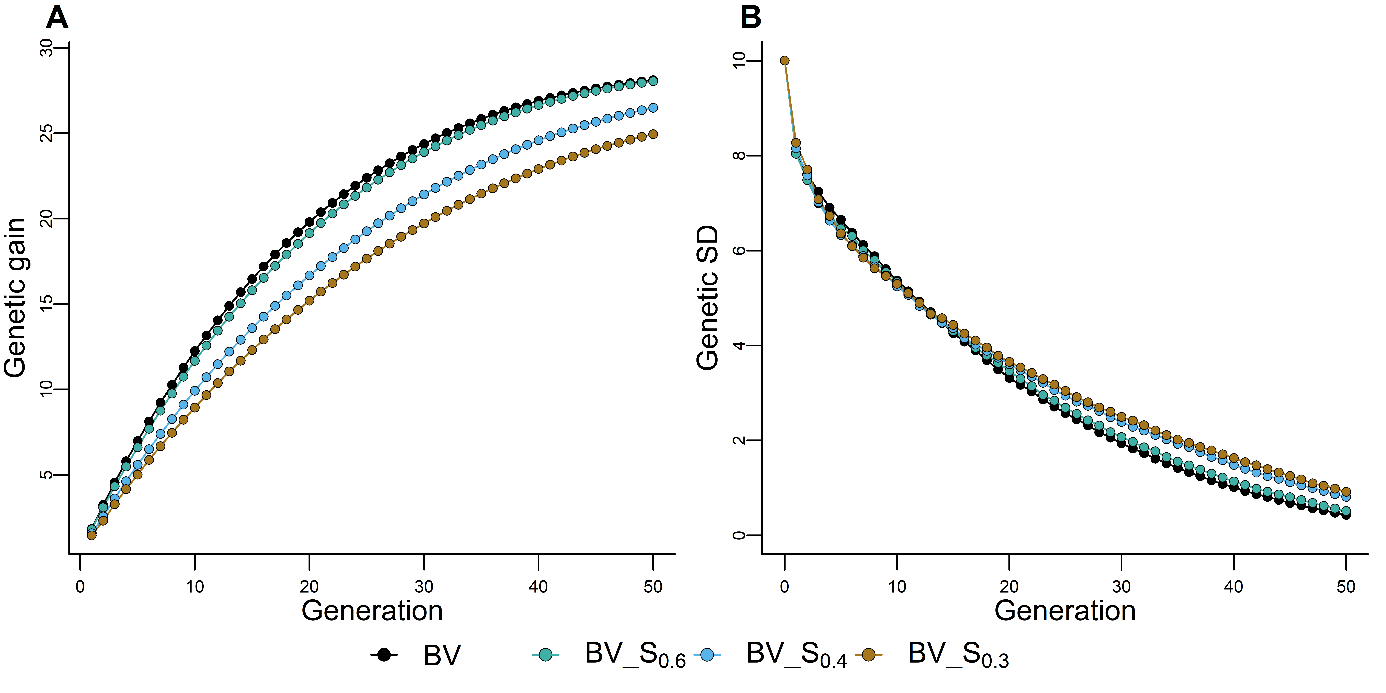

**Figure S6** Effect of similarity matrix (S) on the cumulative genetic gain in genetic standard deviation (A) and genetic standard deviation (B). The selection schemes BV_S_0.6_(_0.4_,_0.3_) optimize mate allocation by maximizing the breeding value (BV) under various constraints (0.6, 0.4, and 0.3) on the haplotype similarity of parents. The results maximizing the index combining breeding value and Mendelian sampling variance are presented in Figure 3. Results are reported for 100 simulation runs.

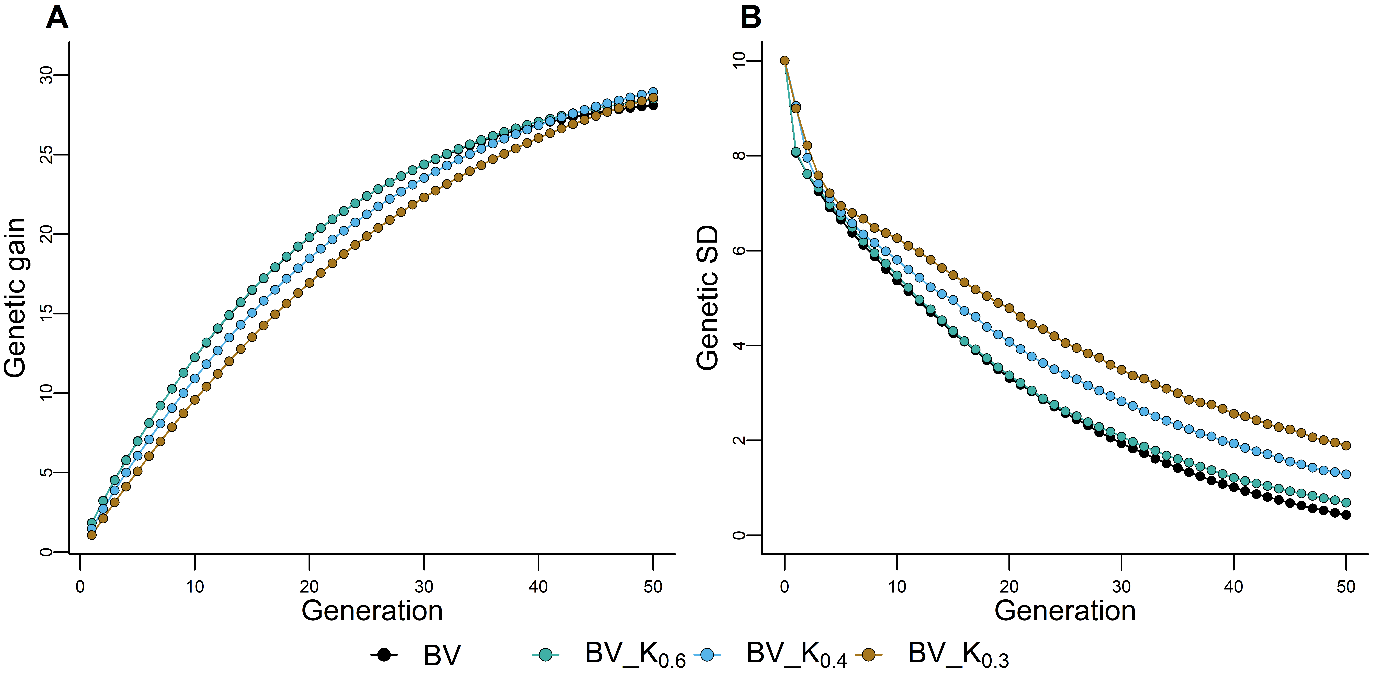

**Figure S7** Effect of similarity standardized matrix (K) on the cumulative genetic gain in genetic standard deviation (A) and genetic standard deviation (B). The selection schemes BV_K_0.6_(_0.4_,_0.3_) optimize mate allocation by maximizing the breeding value (BV) under various constraints (0.6, 0.4, and 0.3) on the standardized haplotype similarity of parents. The results maximizing the index combining breeding value and Mendelian sampling variance are presented in Figure 4. Results are reported for 100 simulation runs.

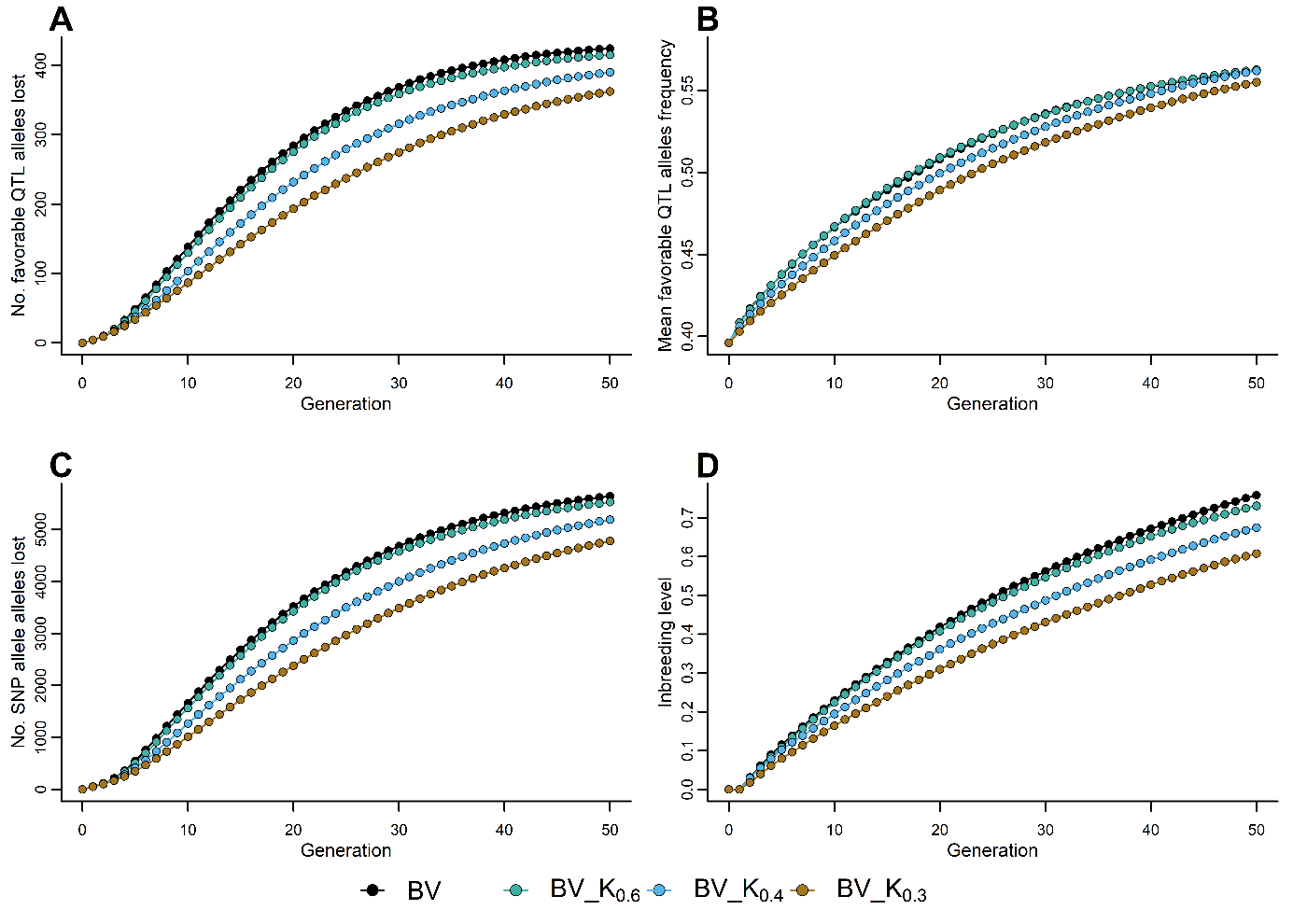

**Figure S8** Effect of standardized similarity matrix on favorable QTL alleles lost (A), mean favorable QTL allele frequency (B), SNPs lost (C), and expected inbreeding rate (D). The selection schemes BV_K_0.6_(_0.4_,_0.3_) optimize mate allocation by maximizing the breeding value (BV) under various constraints (0.6, 0.4, and 0.3) on the standardized haplotype similarity of parents. The results maximizing the index combining BV and Mendelian sampling variance is presented in Figure 5. Results are reported for 100 simulation runs.

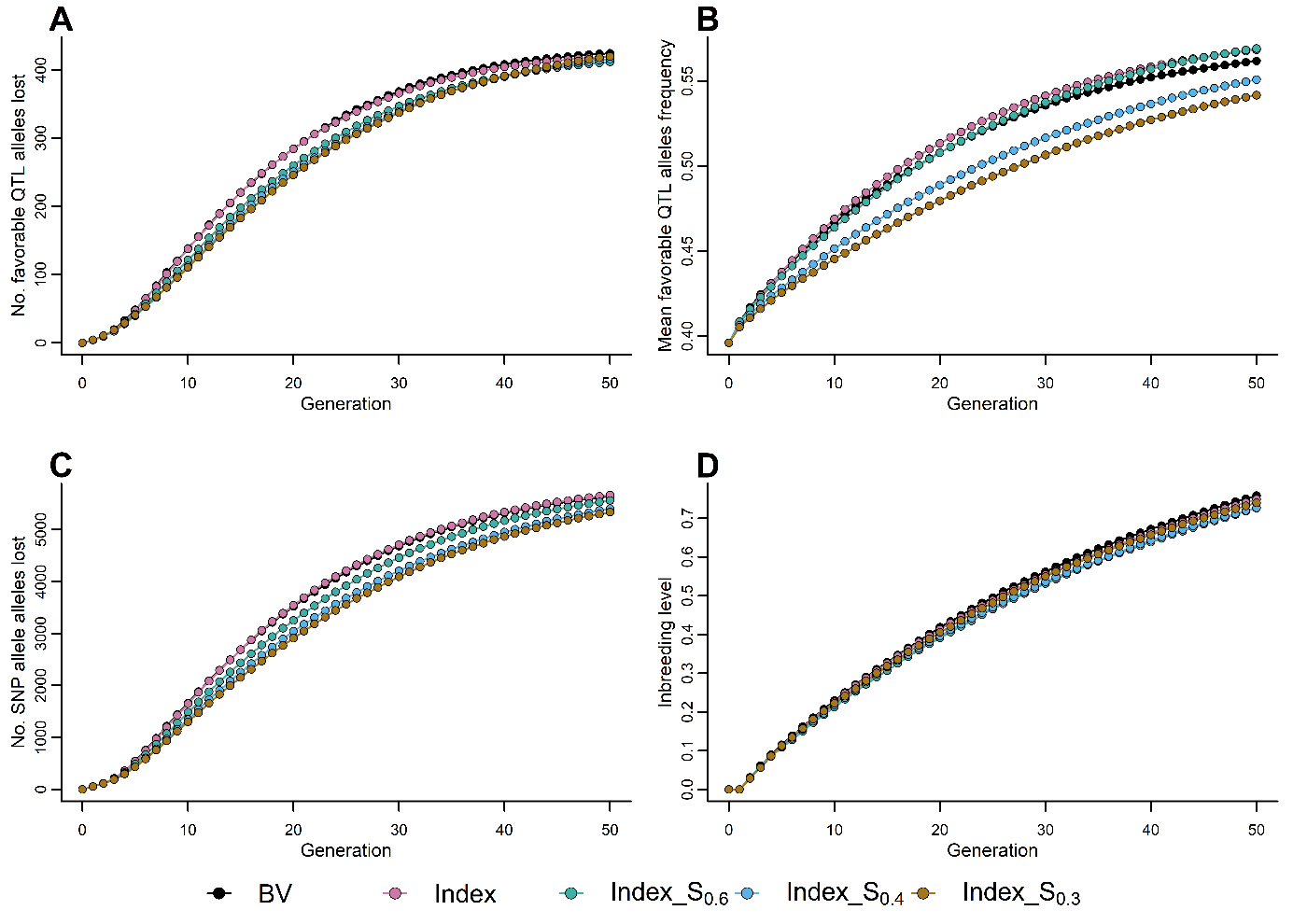

**Figure S9** Effect of similarity matrix on favorable QTL alleles lost (A), mean favorable QTL allele frequency (B), SNPs lost (C), and expected inbreeding rate (D). The selection schemes Index_S_0.6_(_0.4_,_0.3_) optimize mate allocation by maximizing the index combining breeding value (BV) and Mendelian sampling variance under various constraints (0.6, 0.4, and 0.3) on the haplotype similarity of parents. The results maximizing the BV are presented in Figure S10. Results are reported for 100 simulation runs.

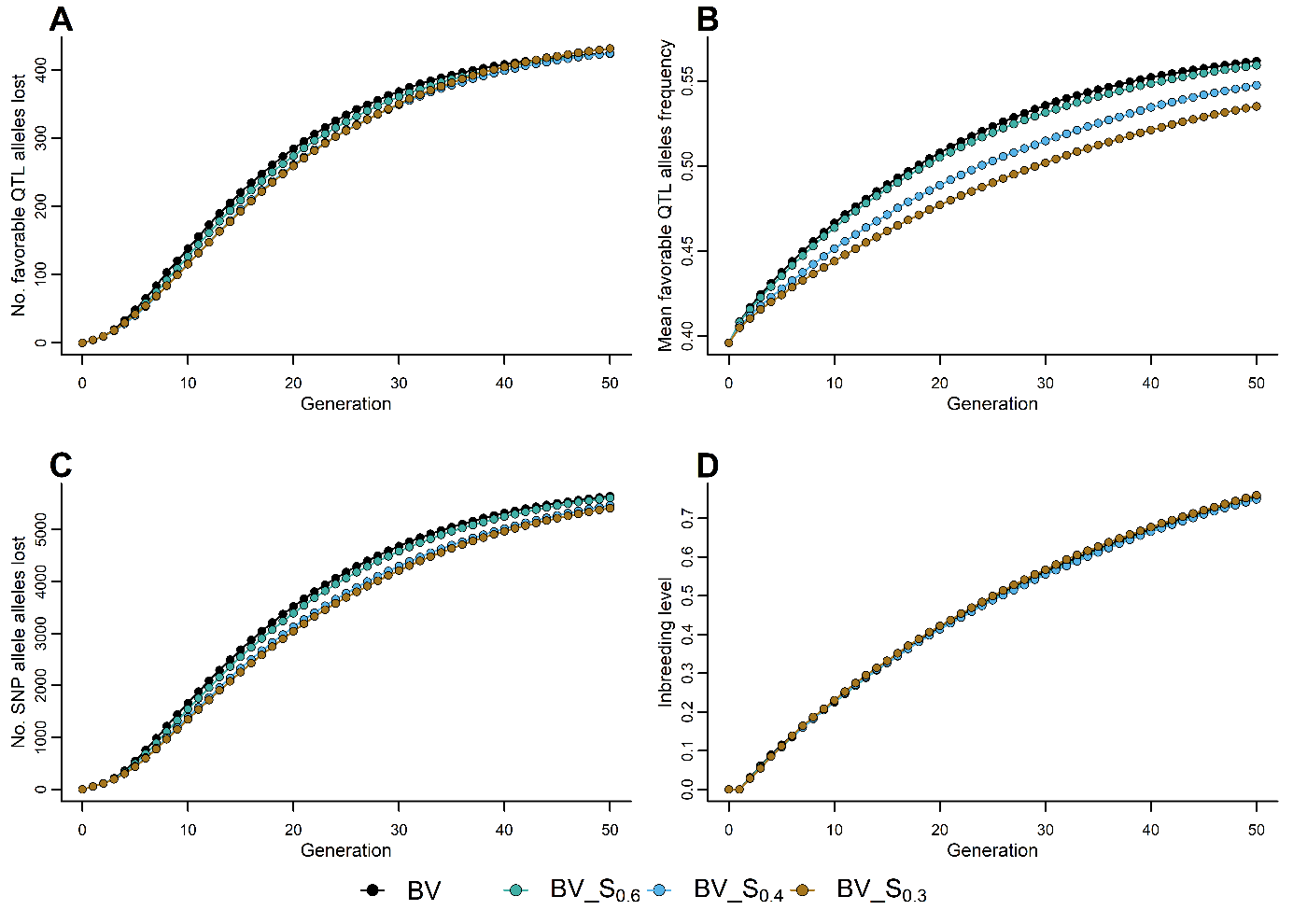

**Figure S10** Effect of similarity matrix on favorable QTL alleles lost (A), mean favorable QTL allele frequency (B), SNPs lost (C), and expected inbreeding rate (D). The selection schemes BV_S_0.6_(_0.4_,_0.3_) optimize mate allocation by maximizing the breeding value (BV) under various constraints (0.6, 0.4, and 0.3) on the haplotype similarity of parents. The results maximizing the BV are presented in Figure S9. Results are reported for 100 simulation runs.

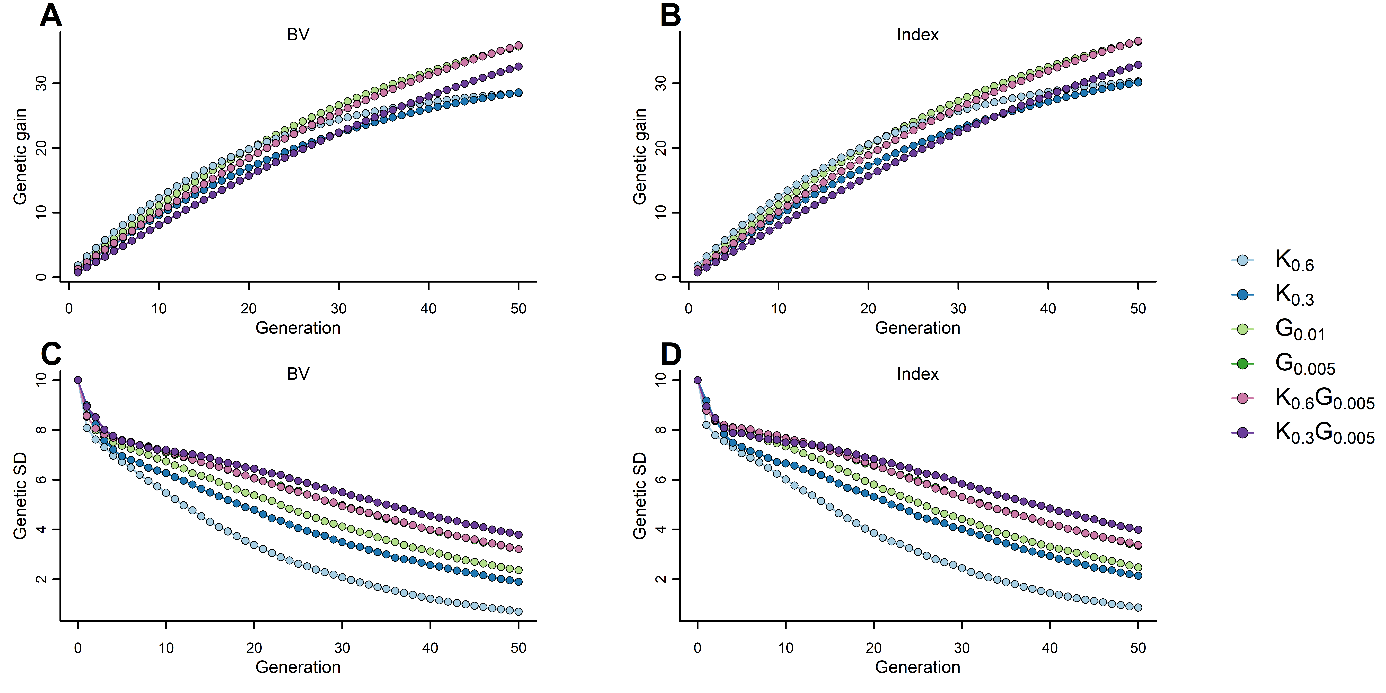

**Figure S11** Effect of standardized similarity matrix (K) and genomic relationship matrix (G) or their combination on the cumulative genetic gain in genetic standard deviation (A) and genetic standard deviation (B) under constraints to select at least five males. The selection schemes optimize mate allocation by maximizing BV (left panels) or index (right panels) within the constraints specified. In the case of K, the constraints are 0.6. 0.4 and 0.3 for standardized haplotype similarity, and 1% or 5% inbreeding rate for G. Results are reported for 100 simulation runs.

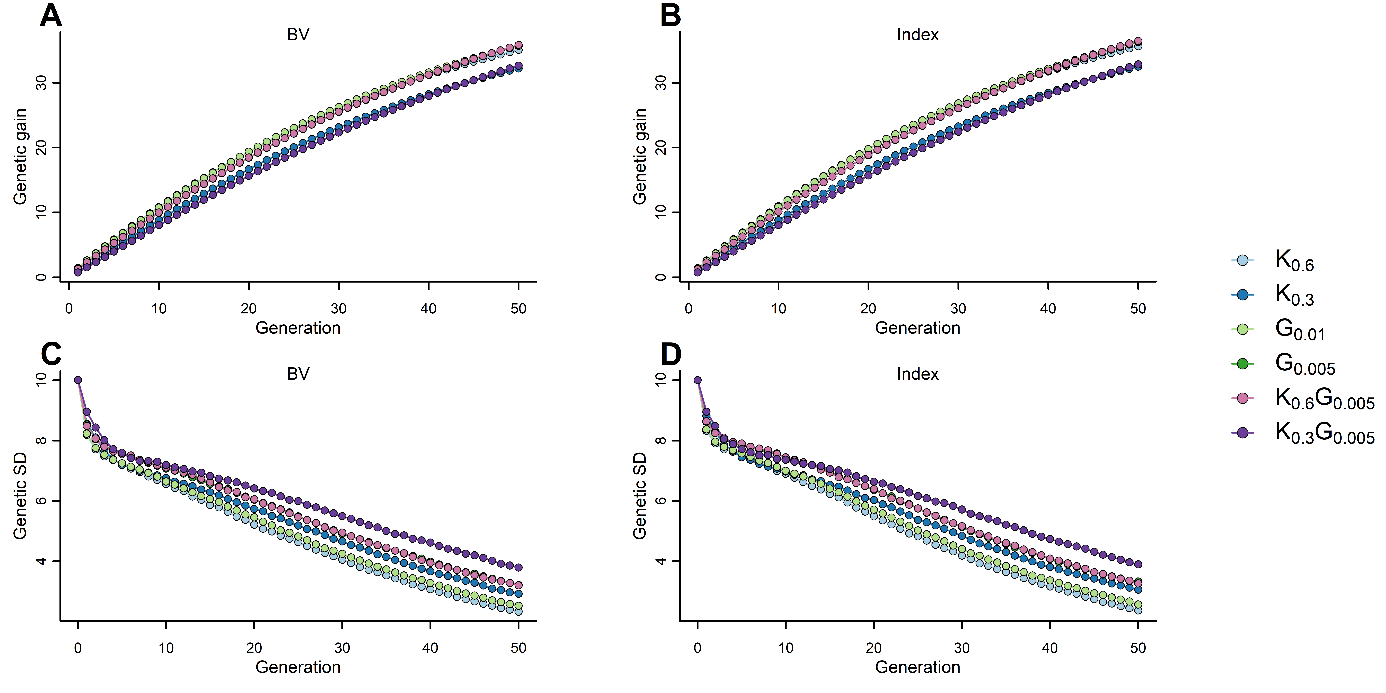

**Figure S12** Effect of standardized similarity matrix (K) and genomic relationship matrix (G) or their combination on the cumulative genetic gain in genetic standard deviation (A) and genetic standard deviation (B) under constraints to select at least 25 males. The selection schemes optimize mate allocation by maximizing BV (left panels) or index (right panels) within the constraints specified. In the case of K, the constraints are 0.6. 0.4 and 0.3 for standardized haplotype similarity, and 1% or 5% inbreeding rate for G. Results are reported for 100 simulation runs.

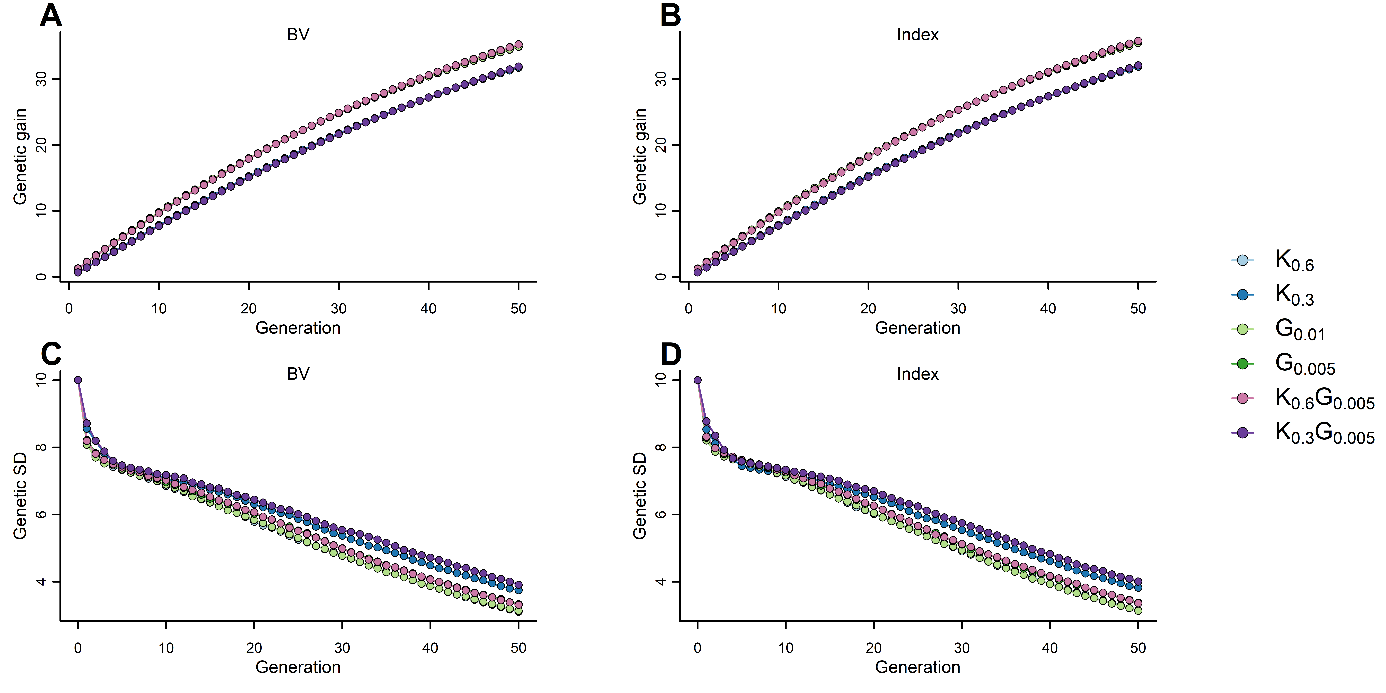

**Figure S13** Effect of standardized similarity matrix (K) and genomic relationship matrix (G) or their combination on the cumulative genetic gain in genetic standard deviation (A) and genetic standard deviation (B) under constraints to select at least 50 males. The selection schemes optimize mate allocation by maximizing BV (left panels) or index (right panels) within the constraints specified. In the case of K, the constraints are 0.6. 0.4 and 0.3 for standardized haplotype similarity, and 1% or 5% inbreeding rate for G. Results are reported for 100 simulation runs.
